# A potent SARS-CoV-2 antibody neutralizes Omicron variant by disassembling the spike trimer

**DOI:** 10.1101/2022.03.21.485243

**Authors:** Wuqiang Zhan, Xiaolong Tian, Xiang Zhang, Shenghui Xing, Wenping Song, Qianying Liu, Aihua Hao, Yuxia Hu, Meng Zhang, Zhenguo Chen, Tianlei Ying, Fei Lan, Lei Sun

**Author notes:** Correspondence: Lei Sun, Fei Lan, Tianlei Ying, and Zhenguo Chen. These authors contributed equally (Wuqiang Zhan, Xiaolong Tian, Xiang Zhang, Shenghui Xing).

## Abstract

The continuous emergence of novel SARS-CoV-2 variants poses new challenges to the fight against the COVID-19 pandemic. The newly emerging Omicron strain caused serious immune escape and raised unprecedented concern all over the world. The development of antibody targeting conserved and universal epitope is urgently needed. A subset neutralizing antibody(nAbs) against COVID-19 from convalescent patients were isolated in our previous study. Here in this study, we investigated the accommodation of these nAbs to SARS-CoV-2 variants of concerns (VOCs), revealing that IgG 553-49 neutralizes pseudovirus of SARS-CoV-2 Omicron variant. In addition, we determined the cryo-EM structure of SARS-CoV-2 spike complexed with three antibodies targeting different epitopes, including 553-49, 553-15 and 553-60. Notably, 553-49 targets a novel conserved epitope and neutralizes virus by disassembling spike trimers. 553-15, an antibody that neutralizes all the other VOCs except omicron, cross-links two spike trimers to form trimer dimer, demonstrating that 553-15 neutralizes virus by steric hindrance and virion aggregation. These findings suggest the potential to develop 49 and other antibody targeting this highly conserved epitope as promising cocktail therapeutics reagent for COVID-19.

**Importance:** The newly emergence of Omicron strain caused higher immune escape, raising unprecedented concerns about the effectiveness of antibody therapies and vaccines. In this study, we identified a SARS-CoV-2 Omicron neutralizing antibody 553-49, which neutralizes Omicron variant by targeting a completely conserved novel epitope. Besides, we revealed that IgG 553-15 neutralizes SARS-CoV-2 by crosslinking virions and 553-60 functions by blocking receptor binding. Comparison of different RBD epitopes revealed that the epitope of 553-49 is hidden in the S trimer and keeps high conservation during SARS-CoV-2 evolution, making 553-49 a promising therapeutics reagent to fight against the emerging Omicron and future variant of SARS-CoV-2.

## Introduction

Coronavirus disease 2019 (COVID-19) caused by the SARS-CoV-2 virus has become a serious pandemic with more than 150 million infections and over 5 million deaths to date(1–3). Coronavirus is enveloped with trimeric transmembrane spike (S) glycoprotein which has essential role in viral entry into host cell. The S1 subunit of S, consisting of receptor binding domain (RBD) and N-terminal domain (NTD), is responsible for receptor binding and S2 subunit primes the fusion of viral and cellular membranes(4). SARS-CoV-2 spike recognizes angiotensin converting enzyme 2(ACE2) of the host cell through receptor binding motif (RBM) on the RBD(5). During infection, spike protein undergoes dynamic conformational change with RBD adopting ‘up’ (or ‘open’) and ‘down’ (or ‘close’) states. Only up RBD can recognize the receptor and initiate the viral entry(4, 6). Spike protein has been used as a major target for development of the vaccine and therapeutic neutralizing monoclonal antibodies (nAbs) against COVID-19(7).

As SARS-CoV-2 escalated to circulate in the human population, many SARS-CoV-2 variants appeared and five of them have been defined as variants of concern (VOCs), including Alpha (B.1.1.7), Beta (P.1), Gamma (B.1.351), Delta (B.1.617.2) and Omicron (B.1.1.529)(8). The spike protein of Alpha strain bears one substitution (N501Y) in the RBM motif, which enhanced its transmissibility(9). The Gamma strain contains three substitutions (K417N, E484K, N501Y) in the RBM motif, resulting in immune escape(10). Delta strain bears L452R, E484Q in the receptor binding domain (RBD) and P681R located in the furin-cleavage site which enhance the fusogenic activity of the spike protein and pathogenicity of the virus(11). Omicron strain, other than K417N, E484A, N501Y, and P681H, contains another 30 mutations in the spike and invalidated almost 85% neutralizing antibodies(12).

Considering the ongoing SARS-CoV-2 pandemic and constantly emerging of variants, antibodies and vaccine targeting the conserved and universal epitopes are urgently needed. Antibodies neutralize virus infection through different mechanisms, such as aggregation of virus particles(13), prevention of viral attachment, inhibition of S fusion or disassembly of virion. The combination of antibodies with different epitopes and various neutralization mechanisms might be an effective way to fight against the SARS-CoV-2 variants. Our previous study identified a set of antibodies from convalescent patients targeting four different epitopes of S, including four RBD-targeting IgGs: 553-15 (abbrev. 15), 553-60 (abbrev. 60), 553-63(abbrev. 63), 553-49(abbrev. 49) and one non-RBD IgG, 413-2(14). Here in this study, we investigated the neutralization activity of these nAbs to SARS-CoV-2 COVs. Binding assay and pseudovirus-based neutralization assay revealed that IgG 60 lost its neutralization ability to all the other VOCs except Alpha. IgG 15 neutralized all the other VOCs except Omicron. Only IgG 49 neutralized SARS-CoV-2 Omicron variant. In addition, cryo-electron microscopy(cryo-EM) structural studies of Omicron S in complex with 49 and wild type (WT) S in complex with 15 and 60, revealed their specific epitopes and neutralization mechanisms. Notably, IgG 49 disassembled Omicron S trimers by targeting a completely conserved novel epitope that is buried inside the trimer and exposed only when the trimers are disassembled. The highly conserved epitope makes IgG 49 and other antibodies targeting this epitope ideal therapeutics reagent to fight against the emerging and future variants of SARS-CoV-2.

## Result

### Broadly neutralizing activity of IgG 49

We first used BLI to investigate the affinity between spike protein of different SARS-CoV-2 variants (WT, Alpha, Beta, Gamma, Delta, Omicron) and five IgGs: 15, 60, 63, 49, and 413-2, to determine whether these monoclonal antibodies accommodate the emerged variants. The Alpha S could bind to all five antibodies, while the Beta and Gamma S proteins were not able to bind with 60, 63 (Fig. S1). IgG 15 retained high binding ability to S of Alpha, Beta, Gamma, Delta at 4.05, 4.25, 9.34 and 2.78nM, respectively, but could not bind Omicron S (Fig. S1). Fortunately, 49 maintained the binding affinity to S of all five VOCs at 0.23, 25.51, 6.99, 7.19 and 15.21 nM, respectively (Fig. 1 and Fig. S1).

**FIG 1.**
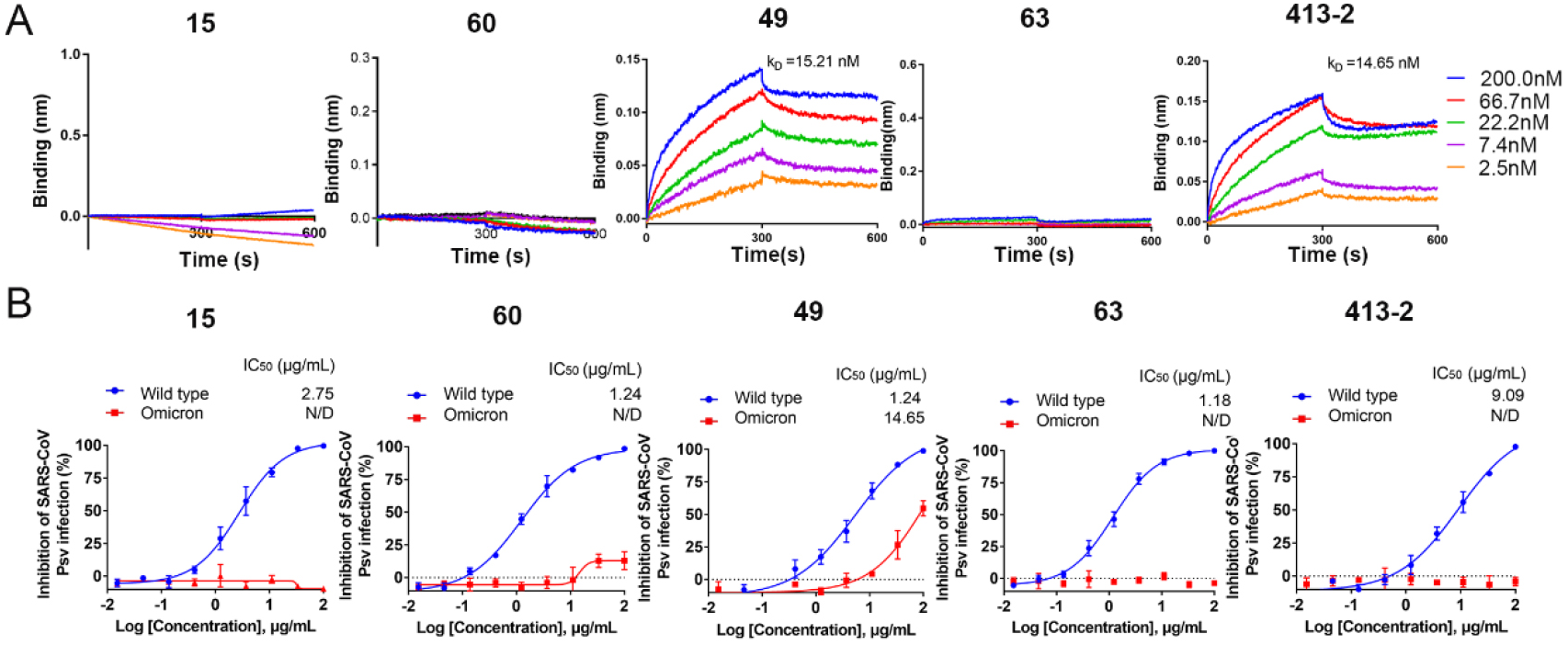
Accommodation of monoclonal antibodies to SARS-CoV-2 Omicron variant. (A) BLI sensorgrams and kinetics of monoclonal antibodies binding to SARS-CoV-2 Omicron spike protein. (B) Neutralizing activity of IgG (15, 60, 49, 63, 413-2) against SARS-CoV-2 WT and Omicron pseudoviruses (PSV).

We next assayed the neutralizing activity of these five antibodies against SARS-CoV-2 variants and SARS-CoV pseudoviruses. They neutralized WT SARS-CoV-2 pseudoviruses with 50% inhibition concentration (IC_50_) values as 2.94, 0.79, 10.82, 0.11, 13.91μg/mL (Fig. S1), respectively. The neutralization activities of 60 and 63 on SARS-CoV-2 Beta and Gamma strains were completely abolished, which is consistent with the result of BLI (Fig. S1). The Delta strain evades the neutralization activity mediated by these antibodies, like 60, 49, 63, 413-2, although the spike protein binds tightly to these antibodies (Fig. S1), which is consistent with the result that Delta pseudovirus is relatively resistant to neutralizing antibodies(15) due to the mutation P681R enhance the fusogenic activity of the spike protein. As to Omicron strain, 49 and 413-2 maintained the binding activity, however only 49 was able to neutralize the pseudovirus with IC_50_ as 14.65ng/mL (Fig.1).

Thus, among the 5 antibodies targeting different epitopes of spike, only IgG 49 retained its affinity with Omicron S and its neutralization activity on Omicron.

### IgG 49 targets conserved epitopes and induces S trimer disassemble

To understand the broad neutralization mechanism of 49, we used Cryo-EM to determine the structure of 49 complexed with prefusion stabilized ectodomain of SARS-CoV2 Omicron S. The purified omicron S trimer was mixed with 49 at 1:1.5 molar ratio, incubated at 4 °C for 2 hour and further purified by gel filtration. The peak fraction of gel-filtration was used for cryo-EM data collection. Two states of particles were observed: Omicron S trimer at apo state (Apo-OS) and S monomer complexed with 49 (OS-49) (Fig.S2, S3, S4). Thus, the Omicron S trimers were disassembled into monomers upon 49 binding. The cryo-EM structure of Omicron S trimer was determined to 3.40 Å (Fig.2C, S3, S4). For S-monomer-49 complex, due to the flexibility of the S monomer, only the RBD-49 region was locally refined to 4.06 Å resolution (Fig.2A, S3, S4), sufficient for model building of RBD and 49(OS-RBD-49) and interface analysis (Fig.2).

**FIG 2.**
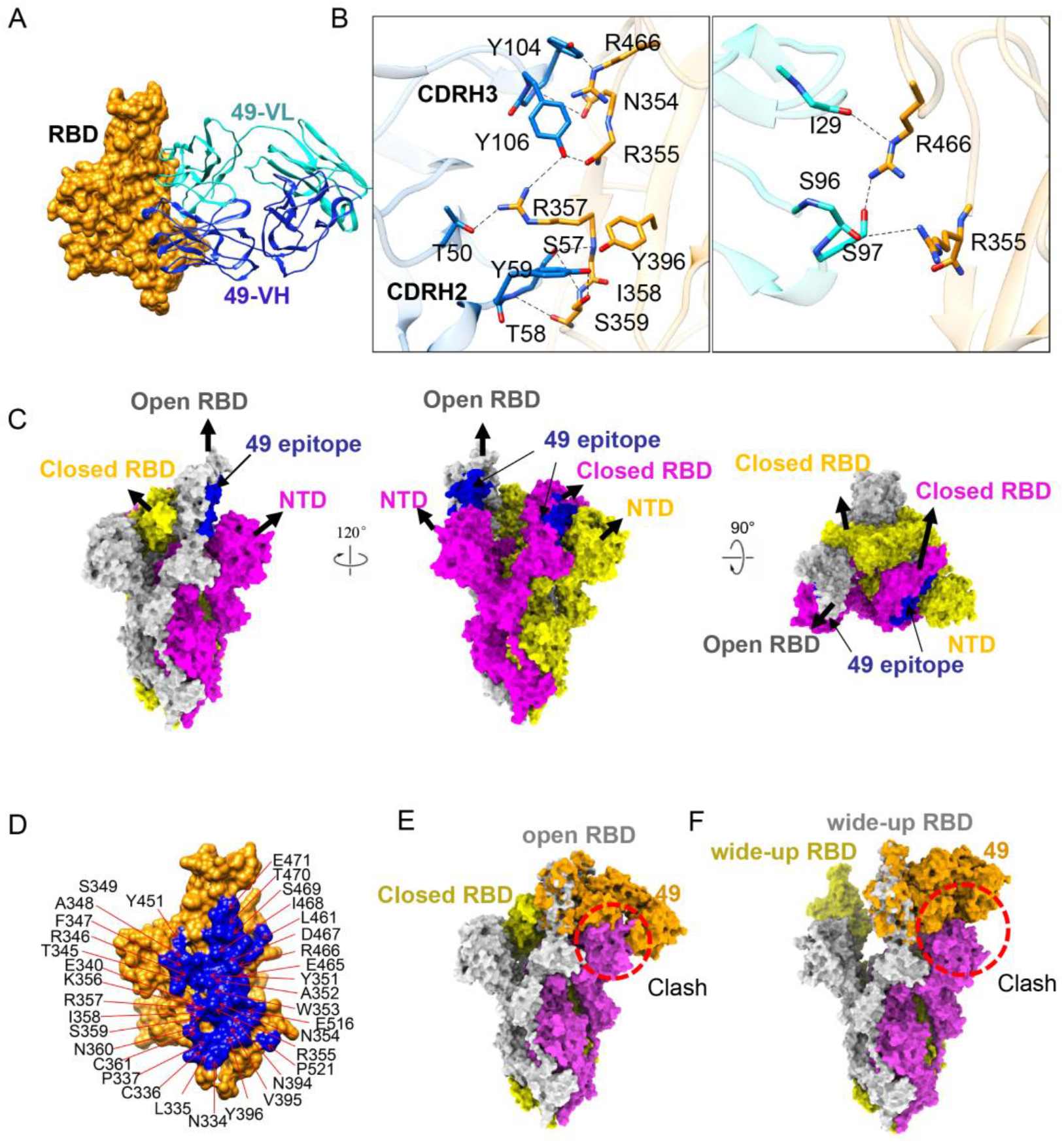
An outer cryptic conserved epitope on SARS-CoV-2 Omicron S RBD recognized by IgG 49. (A) Close-up view of the interactions between 49 and Omicron S-RBD. The RBD is displayed in yellow surface. The heavy chain and light chain of 49 are shown as cartoon colored in blue and cyan, respectively. (B) The interaction of 49-VH(left) and 49-VL(right) with Omicron RBD. (C) The apo Omicron S model displayed in side and top view. The 49 epitope (colored in blue) on Omicron RBD was covered by the NTD of adjacent protomer. (D) Close-up view of 49 epitope. The residues involved in the interaction were labeled. (E-F) Binding of 49 to one RBD-up spike trimer (E) or a wide open RBD (PDB: 7WHK) (F) would clash (indicated by the red dashed circles) with the NTD of adjacent protomer.

IgG49 buries surface area of 1076Å^2^ on RBD (Fig. 2D). The binding of 49 is mainly mediated by CDRH2, CDRH3, CDRL1 and CDRL3 through hydrophilic and hydrophobic interactions (Fig.2A, 2B). Residues N354, R355, R357, I358, S359, Y396, R466 from RBD participate in the interaction by forming 11 pairs hydrogen bonds and 1 pair hydrophobic interaction.

Totally 35 residues of RBD are involved in the interaction and they are completely conserved among all VOCs (Fig.2D, 6), revealing an extremely conservative epitope. In S trimer, this epitope is covered by the NTD of the neighboring protomer no matter the RBD adopts up or down conformation (Fig.2C). Both structural alignment of the OS-RBD-49 complex with apo Omicron S and the wide-open RBD of Omicron S (PDB:7WHK) induced by nanobody(16) binding indicates that 49 would clash with NTD of adjacent protomer (Fig.2E, 2F). The binding of 49 triggers dramatic movement of RBD and NTD, resulting in disassembly of the spike trimer. These findings suggested the potential to develop 49 and other antibody targeting the similar cryptic epitope as promising therapeutics for COVID-19.

### IgG 15 induces the formation of dimer of trimers

To characterize cross-neutralization mechanism of 15, we determined the cryo-EM structure of the prefusion stabilized SARS-CoV-2 S(D614G) ectodomain trimer complexed with 15. Negative stain micrographs showed that incubation of IgG 15 with S crosslinked two S trimers (Fig.S7). Further cryo-EM study unveiled a previously uncharacterized structure of SARS-CoV-2 S trimer in complex with 15, in which three IgG molecules cross-link two S trimers, forming an unsymmetrical head-to-head trimer dimer (Fig.3, S8, S9). The cryo-EM structure of the complex was determined to an overall 4.47 Å (Fig.S8, S9), with RBD-15 region locally refined to 3.80 Å resolutions (Fig.S8, S9). The Fc region is missing in the final reconstruction because of its flexibility, though weak density is observed in the cryo-EM map (Fig.S10).

**FIG 3.**
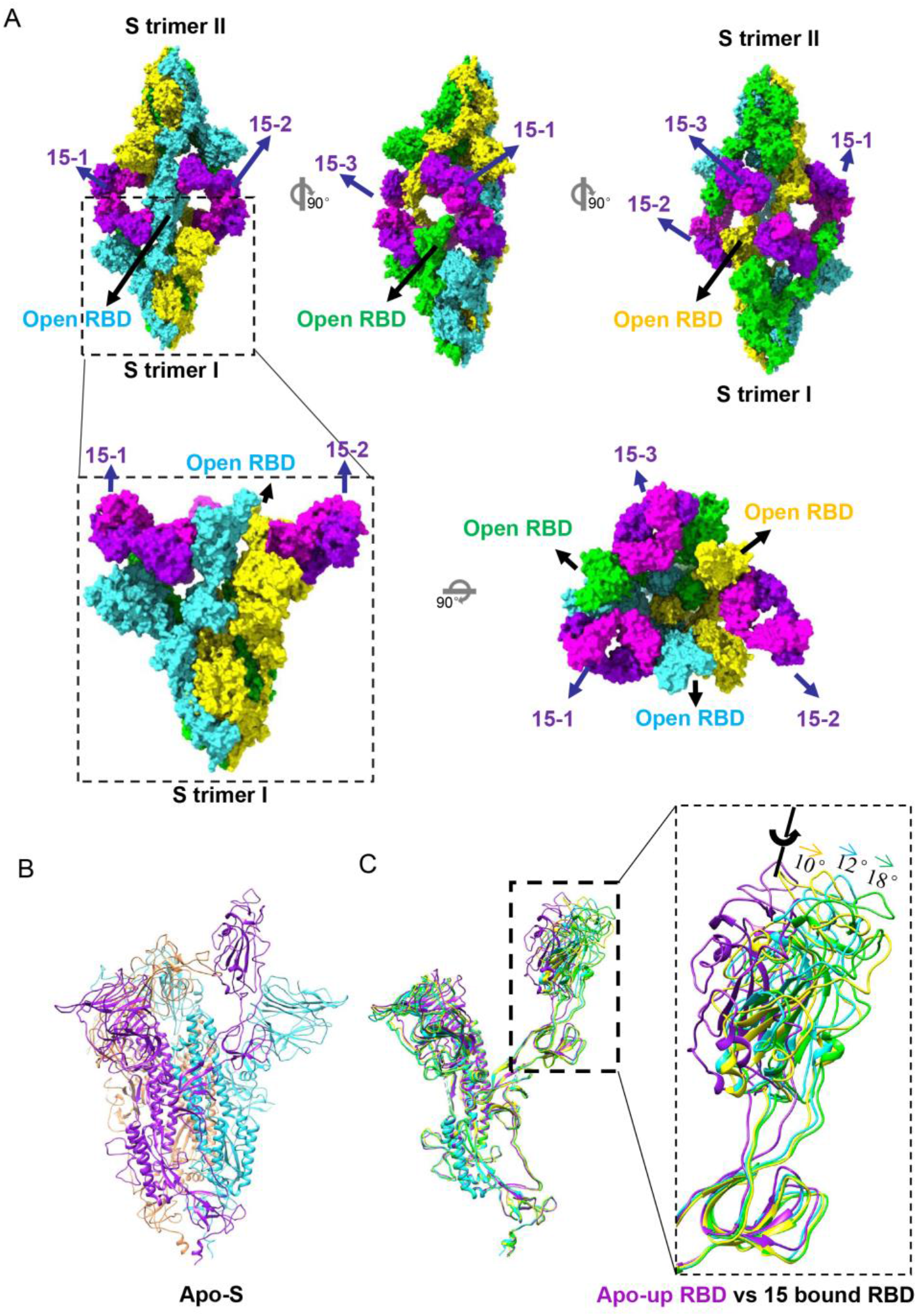
Cryo-EM structures of S D614G in complex with IgG 15. (A) 15 induces spike trimer forming trimer of dimer. Side and top view of the trimer of spike trimer binding three 15 fabs. (B) The model of apo S D614G trimer displayed in ribbon (C) IgG 15 induced all RBDs to open state. Comparison of 15 bound S protomers with apo-up S protomer (medium and right). Purple indicates apo-up S protomer.

Binding of IgG 15 induced all RBDs of S trimers to open state (Fig.3), although the apo-S trimer alone used in this study adopts one “up” RBD and two “down” RBDs (Fig.3B, S5, S6). When compared with the up RBD in the unliganded S trimers, three RBDs rotate by ~18°, ~12° and ~10°, respectively (Fig.3C). Thus, three up RBDs of the S trimer does not follow three-fold symmetry (Fig.3A). Despite the different RBD conformations, all three bound 15 Fabs interact with RBDs in a similar conformation. The interaction is focused on the inner site surface of the RBD, without too much overlapping with the RBM region (Fig.S11). The binding of the 15 Fab with RBD buried 787 Å^2^ surface area. A total of 21 residues from RBD are involved in the interaction, with 9 residues binding to the heavy chain and 17 residues to the light chain (Fig.4B). All three CDRs of heavy chain participate in the binding through hydrophobic and hydrophilic interactions. Extensive hydrophobic contacts are mainly formed between CDRH3 loop (100-YYY-103) and residues Y369, A372, F374, F377, P384 of RBD. Residue A372 inserts into the hydrophobic cavity consist of Y102 of CDRH3, W33 of CDRH1 and Y59 of CDRH2 forming intensive hydrophobic contacts. The 4 pairs hydrogen bonds were formed between Y103 of CDRH3, Q53 of CDRH2 and the loop (370-NSAS-373) of RBD(Fig.3B).

**FIG 4.**
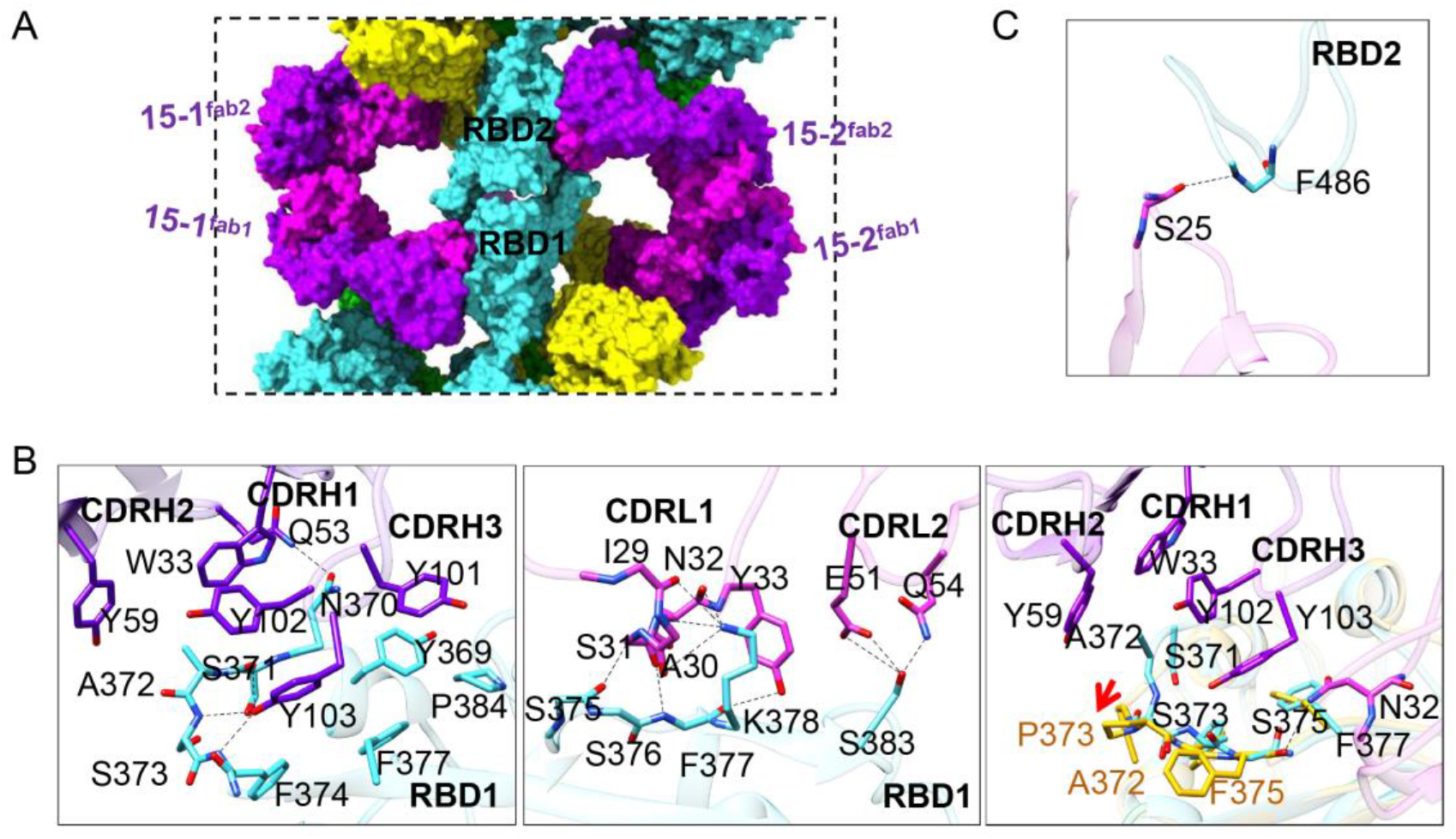
The interaction of 15 and apo S-RBD. (A) Close-up-view of two opposite RBD region. (B) VH of 15 binds the RBD through the hydrophobic and hydrophilic interaction between CDRH3 loop and RBD (369-384) (left); VL of 15 binds the RBD through the CDRL1 and CRL2 loop (medium); The interaction between VH of 15 and Omicron RBD was destroyed by the mutation of S373P (right); (C) VL of 15 binds to the opposite RBD through CDRL1.

The binding of light chain of 15 to RBD is mainly mediated by hydrogen bonds between CDRL1 (29-IASNY-33) and the β sheet (375-STFK-378) of RBD, as well as E51, Q54 of CDRL2 and S383 of RBD. Interestingly, 15-1^fab1^ and 15-2^fab2^ simultaneously interact RBDs of the opposite S protomer, introducing two additional interactions (Fig.3A, 3C). For stabilization of the trimer dimer, VL of 15-1^fab1^ and 15-2^fab2^ binds to the opposite RBD through CDRL1, forming a hydrogen bond between S25 and F486. Therefore, these two patches contact may contribute to the stabilization of the dimer.

Based on the complex structure, the failure of 15 neutralizing the Omicron variant is mainly due to the mutation of S373P which destroyed the hydrophobic contacts mediated by A372 of RBD and the three pairs hydrogen bonds mediated by Y103 of VH (Fig. 3B).

### IgG 15 neutralizes the virus by aggregating virion and and blocking receptor attachment

Although 15 targets the non-RBM region, the previous study showed that 15 was able to compete with ACE2 for binding with membrane-bound S protein(14). Structure superimposition of RBD-ACE2 and S-15 complex indicates that 15 would clash with ACE2 glycosylated residue (N322) (Fig. S11B). A high-throughput study demonstrated that antibodies targeting to the inner site are prone to induce spike-cross linking, causing steric hindrance or virions aggregation to neutralize virus (17). Thus, IgG crosslinking also contribute to the 15 neutralization, as well as antibody targeting to this epitope which is absent of ACE2-blocking.

### IgG 60 epitope overlaps with the RBM region

It has been shown that 60 targets two epitopes of Spike. To identity its precise epitopes, we studied the cryo-EM structure of IgG 60 complex with the stabilized SARS-CoV-2 S ectodomain. Two distinct conformational states were observed (Fig. 5A), corresponding to one up RBD + two down RBDs (1-up) and two up RBDs + one down RBD (2-up). Both structures were determined to 3.25 Å (Fig. S12, S13). In the 1-up state, two 60 molecules bind to the S trimer: one on the up RBD (named as 60-up) and the other one on the down RBD which is clockwise next to the up RBD (named as 60-down). In the 2-up state, each RBD binds to a 60 no matter at up or down conformation (Fig. 5A).

**FIG 5.**
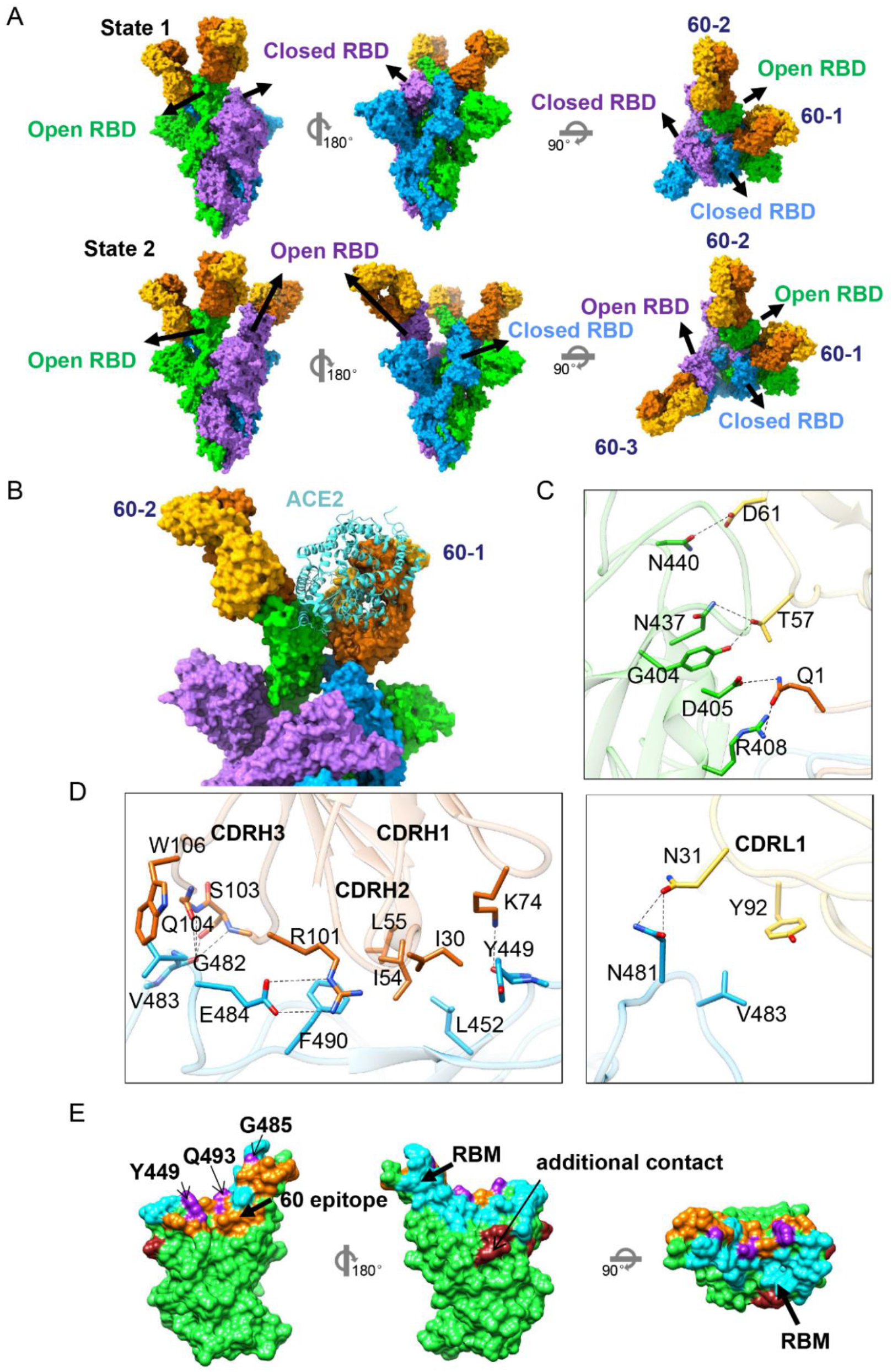
Cryo-EM structures of D614G S in complex with IgG 60. (A) 60 binds to spike trimer as two states. Side and top view of the trimer with three 60 fabs in surface representation. (B) Superimposition of up RBD with RBD-ACE2 (PDB:6LZG) revealed a minor clash between the ACE2 and 60. (C) The interaction between −60-1 and neighboring open RBD. (D) The interface of 60-spike; VH of 60-1 binds the closed RBD through the CDRH1, CDRH2 and CDRH3 loop (left); (C) VL of 60-1 binds the closed RBD through one hydrogen and one patch hydrophobic contacts (right). (E) Surface representation of RBD and buried binding site, including 60 (Orange), 60 additional contact (brown), ACE2 (cyan) and 60 epitope overlaps with RBM region (purple).

**FIG 6.**
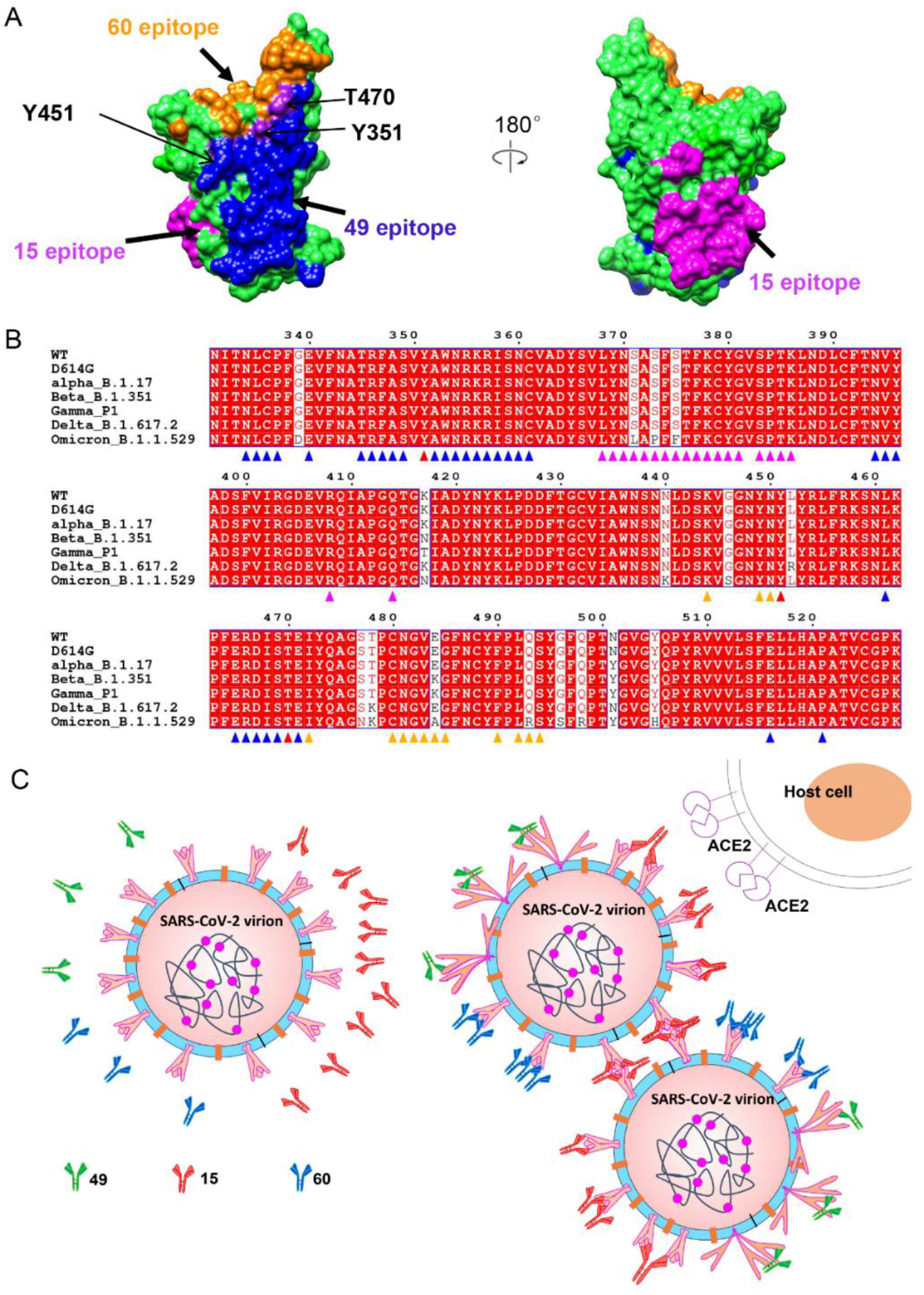
Epitope and neutralization mechanism of three nAbs. (A) Comparison of 49, 15, 60 epitopes on S RBD. S RBD is displayed in green surface. The 49, 15 and 60 epitopes are colored blue, magenta and orange, respectively. (B) Sequence alignment of SARS-CoV-2 WT and all VOCs, showing that IgG 49 targets to a completely conserved epitope. Conserved amino acids are highlighted as red. Residues involved in 49, 15, 60 interactions are marked with triangle in blue, magenta and orange, respectively. Residues involved in both 49 and 60 binding are marked with triangle in red. (C) Schematic model of the neutralization mechanisms of 49, 15 and 60.

The epitope of 60-up is near the RBM region and overlaps with residues G485, Q493 and Y449, which are involved in ACE2 binding (Fig .5E). In contrast to 15, 60 Fab molecule has a smaller footprint, with heavy and light chains burying 557 Å^2^ and 101 Å^2^ surface area, respectively. The interaction between 60 Fab and RBD is primarily driven by CDRH1, CDRH2, CDRH3 and CDRL1 through hydrophilic and hydrophobic interaction (Fig. 5D). The residues G482, E484, Y449, N481 of RBD were involved into the hydrophilic interaction by forming one salt bridge and 7 hydrogen bonds. Hydrophobic contacts formed between V483, F490, L452, Y449 from RBD and Y92, W106, L55, L54, I30 of 60 further enhanced the interaction (Fig. 5D).

In addition, the binding of 60 to the down RBD introduced additional contacts through interaction with the neighboring up RBD (Fig.5B, 5C). The additional contact is primarily due to intensive hydrophilic interaction. Salt bridges and hydrogen bonds are formed between Q1, T57 and D61 from VH and VL and R408, D405, N437, Y508, N440 of open RBD (Fig. 5C). This additional contact locks two S protomers together and hinder RBD further opening. Superimposition of the up RBD with ACE2-SARS-CoV-2 RBD complex structure revealed a minor clash between the ACE2 and 60-up (Fig. 5B). Interestingly, ACE2 clashes with the neighboring 60-down Fab. Thus, the binding of 60 with down RBD not only limits the opening of the RBD, but also prevents the up RBD from recognizing and attaching to ACE2. This explains the mechanism of 60-mediated neutralization. Besides, the cross-linking of S trimer can also be observed from the negative stain of 60-S complex (Figure. S8) even though stable dimers of trimers were not observed in the cryo-EM data. Similar to 15, IgG crosslinking might partially contribute to the antibody neutralization.

## Discussion

The continuous emergence of novel SARS-CoV-2 variants reduced the potency of vaccine and therapeutic antibodies induced by first generation virus. Antibodies targeting conservative epitope are urgently needed to be developed. Here, we identified an outer cryptic conserved epitope of RBD which has never been reported.

Antibodies neutralize virus infection through different mechanisms(18). Many antibodies occupy RBM to block receptor attachment(19, 20). Some antibodies lock the spike in all-RBD down conformation to prevent receptor attachment and conformational change which is essential for viral infection(18, 21). Besides, antibodies may induce spike trimer cross-linking to form steric hindrance or virion aggregation to neutralize virons(17, 22, 23). The neutralization mechanism of antibody disassembling the trimeric spikes was rarely reported, due to the instability of the complex. In this study, we observed that IgG 49 completely disassemble Omicron S trimer into monomers upon binding. It has been reported that bispecific nanobody bn03 induced RBD to a wide-open state and eventually disassembled the spike trimer to neutralize SARS-CoV-2(16). Although occupying different epitopes, both 49 and bn03 would clash with NTD when align to up RBD of apo Omicron S. Thus, the significant movement of RBD and NTD would trigger the disassembling of the spike trimer.

In addition, we determined the structures of SARS-CoV-2 WT S complexed with IgG 553-60 and 553-15. IgG 60 targets the RBM region of S, which explains why it lost its neutralization ability to all the other VOCs except Alpha, since RBM is the most variable region among all variants. IgG 15 neutralizes all the other VOCs except Omicron. IgG 15 targets the inner cryptic surface of RBD, close to the CR3022 site(24), which exhibited highly resistant activity to alpha, beta, gamma and delta, except Omicron. The neutralizing mechanism of antibody targeting to this inner cryptic epitope has remained unclear. For example, COVA1-16(25), only bivalent IgG neutralizes SARS-CoV and SARS-CoV-2 pseudovirus, but not Fab alone. Here, we observed that IgG 15 crosslinks two spike trimers to form trimer dimer, demonstrating its neutralization mechanism by inducing virion aggregation, which may contribute to understand the neutralization mechanism of COVA1-16 antibody in the absence of ACE2 blocking.

Thus, antibodies 49, 15 and 60 neutralize SARS-CoV-2 by different mechanisms. The epitope of 60 partly overlaps with RBM, which is easily exposed on the surface of virus and easily mutated during viral adaptations. IgG 15 targets to a relatively conserved inner site of RBD exposed when at least two RBD are open(24), which enables its neutralizing all the other VOCs except omicron. The epitope of 553-49 is not exposed on the S trimer surface and keeps high conservation during SARS-CoV-2 evolution. Our results provide a new potential therapeutic against SARS-CoV-2 Omicron, and highlight a novel conserved epitope on S RBD that may be used to achieve broad protection against the emerging variants.

## Methods

### Production of recombinant D614G and variants spike ectodomain

The gene encoding SARS-CoV-2 S ectodomain (residues 1-1208, Gene Bank: MN908947) was synthesized (GeneScript) and cloned into mammalian expression constructs pcDNA-3.1, proline substitutions at residues 986 and 987, “GSAS” substitution at furin cleavage site (residues 682-685), a T4 fibritin trimerization motif, an HRV3C protease cleavage site, a TwinStrepTag, and an 8XHisTag at C-terminal were introduced simultaneously by MultiS one step cloning kit (Vazyme). Using the SARS-CoV-2 WT plasmid as the template, mutations such as D614G, B.1.17 (Del 69H70V, N501Y, P681H, S982A, Del 145Y, A570D, T716I, D1118H), B.1.351 (K417N, E484K, N501Y) and P1 (K417T, E484K, N501Y) was introduced by Multi Site-Directed Mutagenesis Kit (Yeasen).

The Human codon gene encoding SARS-CoV-2 Omicron S ectodomain was purchased from GeneScript. The expression plasmid of Omicron S with HexaPro mutations(26), “GSAS” substitution at furin cleavage site (residues 682-285), a T4 fibritin trimerization motif, a TwinStrep Tag, and a C-terminal 8 × His Tag was constructed into pcDNA3.1 vector by MultiS one step cloning kit (Vazyme).

Transiently transfecting suspension HEK293F of the expression plasmid was conducted by polyethylenimine. After 72 hours, the supernatants were harvested and filtered for further purification by Histrap HP (GE) and Superose 6 increase 10/300 column (GE Healthcare) in 20 mM Tris pH 8.0, 200 mM NaCl.

### Expression and Purification of recombinant antibodies

A pair of plasmids separately expressing the heavy- and the light-chain of antibodies were transiently co-transfected into HEK293F cells. After 72 hours, the supernatants with the secretion of antibodies were collected and purified by protein A/G Sepharose (SMART LIFESCIENCES). The bound antibodies on the Sepharose were eluted by 100mM sodium acetate (pH 3.0), and then pH was adjusted to 6.0 by adding 1M Tris (pH 8.5). The purified antibodies were used in following binding and neutralization analyses.

### Biolayer Interferometry (BLI) Binding Assays

BLI was carried out on an OctetRED96 device (Pall FortéBio) to detect the binding kinetics of spike proteins with antibodies. Firstly, the recombinant spike proteins at 20 μg/mL in sodium acetate buffer (pH 5.0) was immobilized 300 seconds onto activated AR2G biosensors (Pall FortéBio), and then immersed in 1-ethyl-3-(3-dimethylaminopropyl) carbodiimide hydrochloride/ N-hydroxysucci-nimide for 300 seconds. Next, the biosensors loaded with saturated spike proteins were incubated with threefold serial dilutions of antibodies with the initial concentration of 300 nM at 37 °C for 300 seconds, and then immersed into wells containing the kinetics buffer of 0.02% PBST (buffer supplemented with 0.02% Tween 20) for another 300 s. Baseline was established in the kinetics buffer. For data analysis, all the curves were fitted by a 1:1 binding model using the Data Analysis software (FortéBio). Mean kon, koff, and KD values were determined by averaging binding curves within a dilution series having R2 values of greater than 95% confidence level.

### Pseudotyped Virus Neutralization

The pseudotyped virus neutralization assay was performed to determine the neutralization activity of antibodies involved in this study. Briefly, whole spike glycoprotein sequences of SARS-CoV, wild type or variants of SARS-CoV-2 were inserted into the vector of pcDNA3.1+ and severally co-transfected into 293 T cells (ATCC, Manassas, VA, USA) with a defective HIV-1 genome that encodes luciferase reporter. Supernatants containing spike pseudoviruses were collected 48 hours post-transfection. Serial 1/3 dilutions of antibodies were incubated with pseudoviruses at 37°C for 1 hour, and then the mixtures were added in ACE2 expressed Huh-7 cells (104 per well in 96-well plates). All cell lines were cultured in Dulbecco’s modified Eagle’s medium (DMEM) with 10% fetal bovine serum (FBS). Culture medium was refreshed 12 hours post-infection and followed by an additional 48 hours incubation. Huh-7 cells were subsequently lysing with 50 μL lysis reagent (Promega), and 30 μL of the lysates were transferred to 96-well Costar flat-bottom luminometer plates (Corning Costar) for the detection of relative light units using the Firefly Luciferase Assay Kit (Promega) on the instrument of an Ultra luminometer (Tecan). A nonlinear regression analysis was performed on the resulting curves using Prism (GraphPad) to calculate half-maximal inhibitory concentration (IC_50_) values.

### The formation of SARS-CoV-2 S-15 and SARS-CoV-2 S-60 complex

The S trimer at 1.032 mg/mL were mixed with 60 or 15 at 1 mg/mL in a 1:1.4 molar ratio (S trimer: IgG), incubated at 4 °C for 1 hour and further purified by Superose 6 increase 10/300 column (GE Healthcare). The peak tube was concentrated to 0.4-0.5 mg/mL in 20 mM Tris pH8.0, 200 mM NaCl.

### The formation of SARS-CoV-2 Omicron S-49 complex

The omicron S trimer at 1.928 mg/mL were mixed with 49 1.549 mg/mL in a 1:1.5 molar ratio (Omicron S trimer: IgG), incubated at 4 °C for 2 hour and further purified by Superose 6 increase 10/300 column (GE Healthcare). The peak tube was concentrated to 0.4-0.5 mg/mL in 20 mM Tris pH8.0, 200 mM NaCl.

### Cryo-EM sample preparation

3uL complex sample was added to a freshly glow-discharged holey amorphous nickel-titanium alloy film supported by 400 mesh gold grids. The sample was plunged freezing in liquid ethane using Vitrobot IV (FEI/Thermo Fisher Scientific), with 2 s blot time and −3 blot force and 10 s wait time.

### Cryo-EM data collection

Cryo-EM data were collected on a Titan Krios microscope (Thermo Fisher) operated at 300 kV, equipped with a K3 summit direct detector (Gatan) and a GIF quantum energy filter (Gatan) set to a slit width of 20 eV. Automated data acquisition was carried out with SerialEM software(27).

For S-15 complex, movies were taken in the super-resolution mode at a nominal magnification 81,000×, corresponding to a physical pixel size of 1.07 Å, and a defocus range from −1.2 to −2.5 μm. Each movie stack was dose-fractionated to 41 frames with a total exposure dose of about 61 e−/Å^2^.

For S-60 complex, movies were taken in the super-resolution mode at a nominal magnification 105,000×, corresponding to a physical pixel size of 0.82 Å, and a defocus range from −1.2 to −2.5 μm. Each movie stack was dose-fractionated to 40 frames with a total exposure dose of about 60 e−/Å^2^.

For Omicron S-49 complex, movies were taken in the super-resolution mode at a nominal magnification 81,000×, corresponding to a physical pixel size of 1.064 Å, and a defocus range from −1.2 to −2.5 μm. Each movie stack was dose-fractionated to 40 frames with a total exposure dose of about 58 e−/Å^2^.

### Cryo-EM image processing

All the data processing was carried out using either modules on, or through, RELION v3.0 and cryoSPARC(28).

For S-15 complex (Fig.S10), a total of 6,943 movie stacks was binned 2 × 2, dose weighted, and motion corrected using MotionCor2(29). Parameters of contrast transfer function (CTF) were estimated by using Gctf(30). All micrographs then were manually selected for further particle picking upon ice condition, defocus range and estimated resolution. Particles were initially auto-picked by using the Laplacian-of-Gaussian method and then subjected into two-dimensional (2D) classification. The top-class averages were used as the 2D reference for template-picking, yielding 403,685 particles. Binned 4 × 4 particles (4.28 Å/pixel) were extracted and subjected to a routine process of 2D classification, 3D initial model, 3D classification and 3D auto-refine followed by subsequent re-centering and re-extraction binned 1 × 1 (1.07 Å/pixel). Finally, 188,942 particles were grouped and subjected to auto refinement and postprocessing, yielding a trimer dimer density map at 4.47-Å overall resolution with C1 symmetry.

To get a higher-resolution asymmetric single-trimer map, particles were subjected to symmetry expansion with C2 symmetry. All 377,884 expanded-symmetry particles were auto-refined, CTF-refined and polished, yielding the trimer map at 3.45-Å overall resolution with C1 symmetry. To improve resolution at the three RBD-15 interfaces of a trimer, volumes were erased in Chimera(31) and the regions corresponding to the NTD/RBD domains with Fab (NRAb) were used to generate respective local mask (4-pixel extension, 8-pixel soft cosine edge). Three copy of particles targeted on different NRAb regions (NRAb1, NRAb2, NRAb3) were subjected to focused 3D classification without alignment (tau_fudge = 40) by using the local mask separately. 169,450 of NRAb1 particles were selected and then subtracted for focused refinement, yielding a 3.89-Å overall resolution density map. 199,441 of NRAb2 particles were selected and then subtracted for focused refinement, yielding a 3.80-Å overall resolution density map. 249,722 of NRAb3 particles were selected and then subtracted for focused refinement, yielding a 4.11-Å overall resolution density map. The focused refined maps of NRAbs were fitted into map of trimer and merged with it using “vop maximum” command in UCSF Chimera. The composite map for S-15 asymmetric trimer dimer was also formed by using “vop maximum” with 2 × NRAbs maps and 2 × trimer maps, which were all fitted in original 4.47-Å trimer dimer map.

For S-60 (Fig.S13), a total of 3515 movie stacks was motion-corrected and CTF-estimated. 3450 among these micrographs were selected for further processing. After template-picking by 2D reference generated with particles of Laplacian-of-Gaussian picking, 789,443 particles were extracted binned 4 × 4 (3.28 Å/pixel) and subjected to a routine process of 2D classification, 3D initial model, 3D classification and 3D auto-refine followed by subsequent re-centering and re-extraction binned 1 × 1 (0.82 Å/pixel). After 3D classification, two dominant classes were selected separately. 88,701 particles of Class_2 were reconstructed into a 3.25-Å overall resolution density map with two RBD domains down and one up. 99,762 particles of Class_3 were also reconstructed into a 3.25-Å overall resolution density map but with one RBD domain down and two up. To improve resolution at the RBD-mAb interfaces of SmAb60 protein, the same procedure was carried out. Three cluster of particles targeted on different NRAb regions (NRAb1 from Class_2, NRAb2 from Class_2, NRAb3 from Class_3) were subjected to focused 3D classification without alignment (tau_fudge = 40) by using the local mask separately. 88,701 of NRAb1 particles were selected and then subtracted for focused refinement, yielding a 3.35-Å overall resolution density map. 57,260 of NRAb2 particles were selected and then subtracted for focused refinement, yielding a 3.79-Å overall resolution density map. 29,291 of NRAb3 particles were selected and then subtracted for focused refinement, yielding a 3.94-Å overall resolution density map. The focused refined maps of NRAb1/NRAb2 were fitted in map of Class_2 and merged with it using “vop maximum” command in UCSF Chimera. Similarly, the focused refined maps of NRAb1/NRAb2/NRAb3 were fitted in map of Class_3 and merged with it using “vop maximum” command in UCSF Chimera.

For Apo S (D614G) (Fig.S6), routine procedure was carried out. At last, 2,001.157 particles yielded a 2.70 −Å map of apo spike.

For Omicron S-49 (Fig.S4), 7,294 movies were motion-corrected and CTF-estimated. Among these, 5,896 were selected for further processing. After blob-picking in cryoSPARC and 2D classification, trimer and monomer particles were observed. Separate hetero refinements resulted into two major structures: Omicron S trimer at apo state and spike monomer complexed with 49. Finally, 28,589 trimer particles yielded a 3.40-Å map of apo Omicron spike after NU-refinement. And 597,461 monomer particles yielded a 4.06-Å map of RBD-49 interface region.

The reported resolutions above are based on the gold-standard Fourier shell correlation (FSC) 0.143 criterion. All the visualization and evaluation of 3D density maps were performed with UCSF Chimera(31), and the local resolution variations were calculated using RELION(32). These composite maps were then “vop zflip” to get the correct handedness in UCSF Chimera and used for subsequent model building and analysis.

### Model building and structure refinement

Apo-OS trimer model were generated by swiss-model and fitted into the map using UCSF Chimera(31). The model was manually adjusted and glycans at N-linked glycosylation sites were added in COOT(33), then several iterative rounds of real-space refinement were carried out in PHENIX(34).

For Omicron S complexed with 49, the S-RBD model obtained from the apo-OS trimer model and the antibody model generated by swiss-model were fitted into the map using UCSF Chimera(31). The model was further refined in COOT(33) and PHENIX(34) iteratively.

For model building of S D614G (apo-S) trimer model, the WT S model (PDB: 6VYB) were fitted into the map using UCSF Chimera(31) and further refined in COOT(33) and PHENIX(34) iteratively.

For model building of S-15 and S-60, the apo-S trimer model and the antibody (15 and 60) model generated by swiss-model were fitted into the map using UCSF Chimera(31) and followed by manually adjustment in COOT(33), as well as real space refinement in Phenix. The RBD-15 fab (15-S NR15) region was built and refined against the locally refined map (NRAb2) and then docked back into the global refinement trimer maps. The RBD-60 fab (60-S NR60) region was built and refined against the locally refined map (NRAb1) and then docked back into the global refinement trimer maps. Model validation was performed using MolProbity. Details of the refinement statistics of the complexes are summarized in Table S1. Figures were prepared using UCSF Chimera and UCSF ChimeraX(35).

## Acknowledgements

We thank Center of Cryo-Electron Microscopy, Fudan University and Center for Biological Imaging of Institute of Biophysics (IBP) for the supports on cryo-EM data collection. This work was supported by a grant from the major project of Study on Pathogenesis and Epidemic Prevention Technology System (2021YFC2302500) by the Ministry of Science and Technology of China (L.S.).

## Author contributions

L.S., F.L., T.Y and Z.C. conceived and supervised the study. W.Z., S.X., Q.L., A.H. and Y.H. purified the spike proteins and IgG antibodies. X.Z. and Z.C. performed cryo-EM study. X.T and W.S. performed BLI assays and pseudovirus neutralization assay. W.Z., X.Z., Z.C. and L.S. analyzed the data. The manuscript was written by W.Z, L.S. and reviewed, commented and approved by all the authors.

## Data Availability

Coordinates and maps associated with data reported in this manuscript will be deposited to the Electron Microscopy Data Bank (EMDB) and Protein Data Bank (PDB) with accession numbers EMD-32638 and PDB 7WO4 (S-15 trimer dimer), EMD-32639 and PDB 7WO5 (S-15 trimer), EMD-32641 and PDB 7WO7 (S-15 NRAb2 local refinement), EMD-32646 and PDB 7WO8 (S-60 trimer with 2-up state RBD binding 2 fabs), EMD-32647 and PDB 7WOB (S-60 trimer with 3-up state RBD binding 3 fabs), EMD-32648 and PDB 7WOC (S-60 NRAb1 local refinement), EMD-32651 and PDB 7WOG (Omicron RBD-49 Fab), EMD-32901 and PDB 7WZ1 (Apo Omicron S), EMD-32902 and PDB 7WZ2 (Apo S D614G).

---

**TABLE S1.** Cryo-EM data collection and refinement statistics.

|  | OS<br>Trimer | OS_<br>RBD-49 | Apo-S<br>D614G<br>Trimer | S-15<br>trimer<br>dimer | S-15<br>trimer | S-15<br>NR15 | S-60<br>2-up<br>trimer | S-60<br>3-up<br>trimer | S-60<br>NR60 |
| --- | --- | --- | --- | --- | --- | --- | --- | --- | --- |
| **Data collection and processing** |  |  |  |  |  |  |  |  |  |
| Magnification | 81,000 |  | 81000 |  | 81,000 |  |  | 105,000 |  |
| Voltage (kV) | 300 |  | 300 |  | 300 |  |  | 300 |  |
| Total dose (e-/Å<sup>2</sup>) | 58 |  | 58 |  | 61 |  |  | 60 |  |
| Defocus range (μm) | -1.2 to -2.5 |  | -1.2 to -2.5 |  | -1.2 to -2.5 |  |  | -1.2 to -2.5 |  |
| Pixel size (Å) | 1.064 |  | 0.82 |  | 1.07 |  |  | 0.82 |  |
| Symmetry imposed | C1 |  | C1 |  | C1 |  |  | C1 |  |
| Final particles (no.) | 128,589 | 597,461 | 2,001,157 | 188,942 | 377,884 | 199,441 | 88,701 | 99,762 | 88,701 |
| Map resolution (Å) | 3.40 | 4.06 | 2.70 | 4.47 | 3.45 | 3.80 | 3.25 | 3.25 | 3.35 |
| **Refinement** |  |  |  |  |  |  |  |  |  |
| **R.m.s. deviations** |  |  |  |  |  |  |  |  |  |
| Bond lengths (Å) | 0.002 | 0.005 | 0.003 | 0.002 | 0.002 | 0.003 | 0.002 | 0.002 | 0.002 |
| Bond angles (°) | 0.615 | 0.821 | 0.680 | 0.437 | 0.535 | 0.695 | 0.406 | 0.429 | 0.458 |
| **Validation** |  |  |  |  |  |  |  |  |  |
| MolProbity score | 2.45 | 3.15 | 2.35 | 2.17 | 2.23 | 2.64 | 2.10 | 2.24 | 2.29 |
| Clashscore | 7.53 | 14.68 | 6.44 | 5.82 | 5.80 | 6.92 | 6.56 | 7.05 | 6.74 |
| Rotamer outlier (%) | 5.36 | 9.35 | 5.57 | 3.89 | 4.31 | 8.82 | 4.23 | 4.57 | 4.60 |
| **Ramachandran plot** |  |  |  |  |  |  |  |  |  |
| Favored (%) | 92.17 | 80.06 | 93.41 | 93.93 | 93.46 | 90.29 | 96.19 | 95.01 | 93.67 |
| Allowed (%) | 7.83 | 18.52 | 6.59 | 6.06 | 6.54 | 9.71 | 3.81 | 4.99 | 6.33 |
| Disallowed (%) | 0.00 | 1.41 | 0.00 | 0.01 | 0.00 | 0.00 | 0.00 | 0.00 | 0.00 |
| **EMDB** | 32901 | 32651 | 32902 | 32638 | 32639 | 32641 | 32646 | 32647 | 32648 |
| **PDB** | 7WZ1 | 7WOG | 7WZ2 | 7WO4 | 7WO5 | 7WO7 | 7WOA | 7WOB | 7WOC |

**FIG S1.**
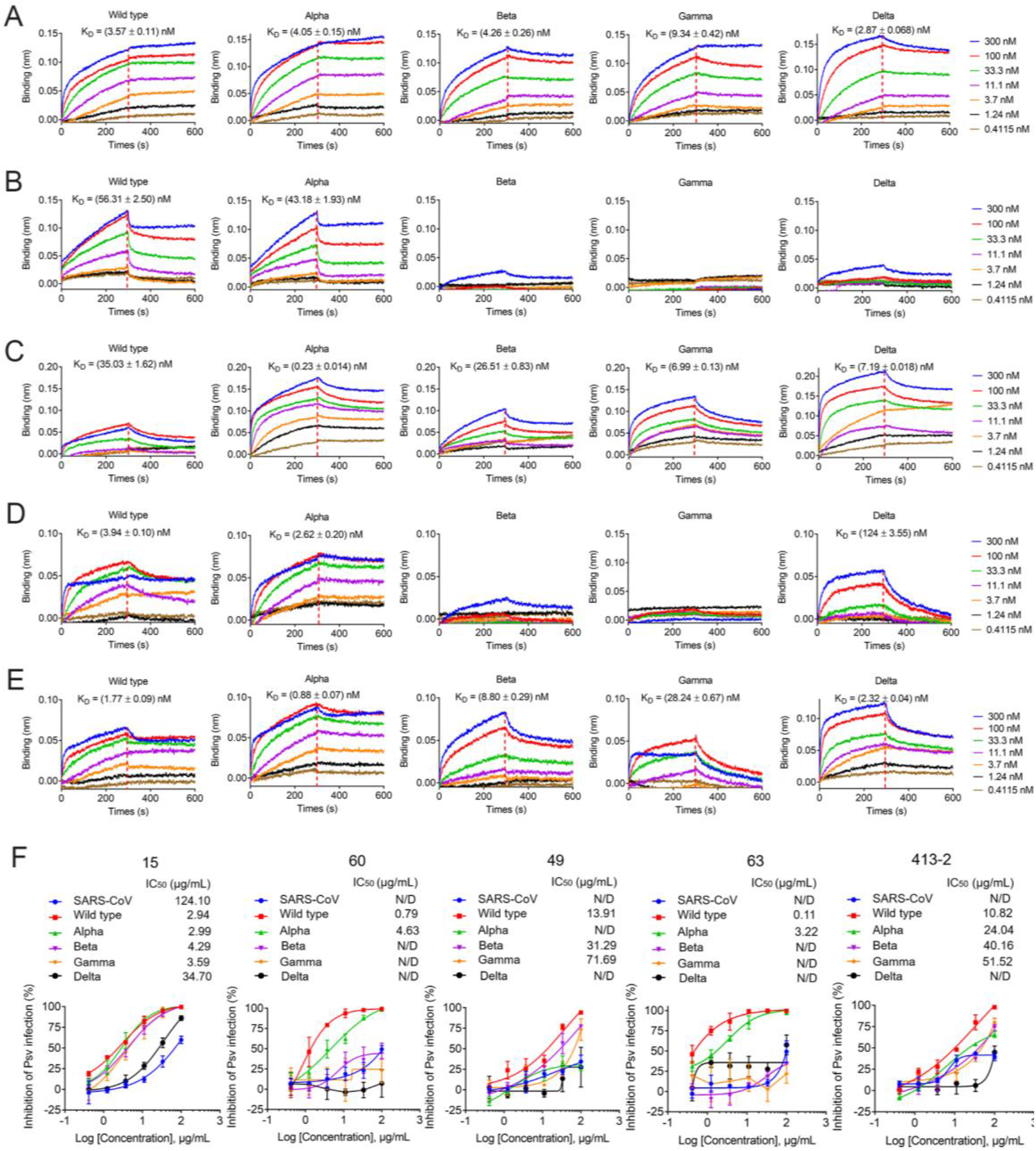
Accommodation of monoclonal antibodies to SARS-CoV-2 VOCs. (A-E) BLI sensorgrams and kinetics of monoclonal antibodies binding to spike protein of SARS-CoV-2 Variants. 15(A) 60(B) 49(C) 63(D) 413-2(E). (F) Neutralizing activity of IgG against SARS-CoV-2, SARS-CoV-2 variants (Alpha, Beta, Gamma, Delta) and SARS-CoV pseudoviruses (PSV).

**FIG S2.**
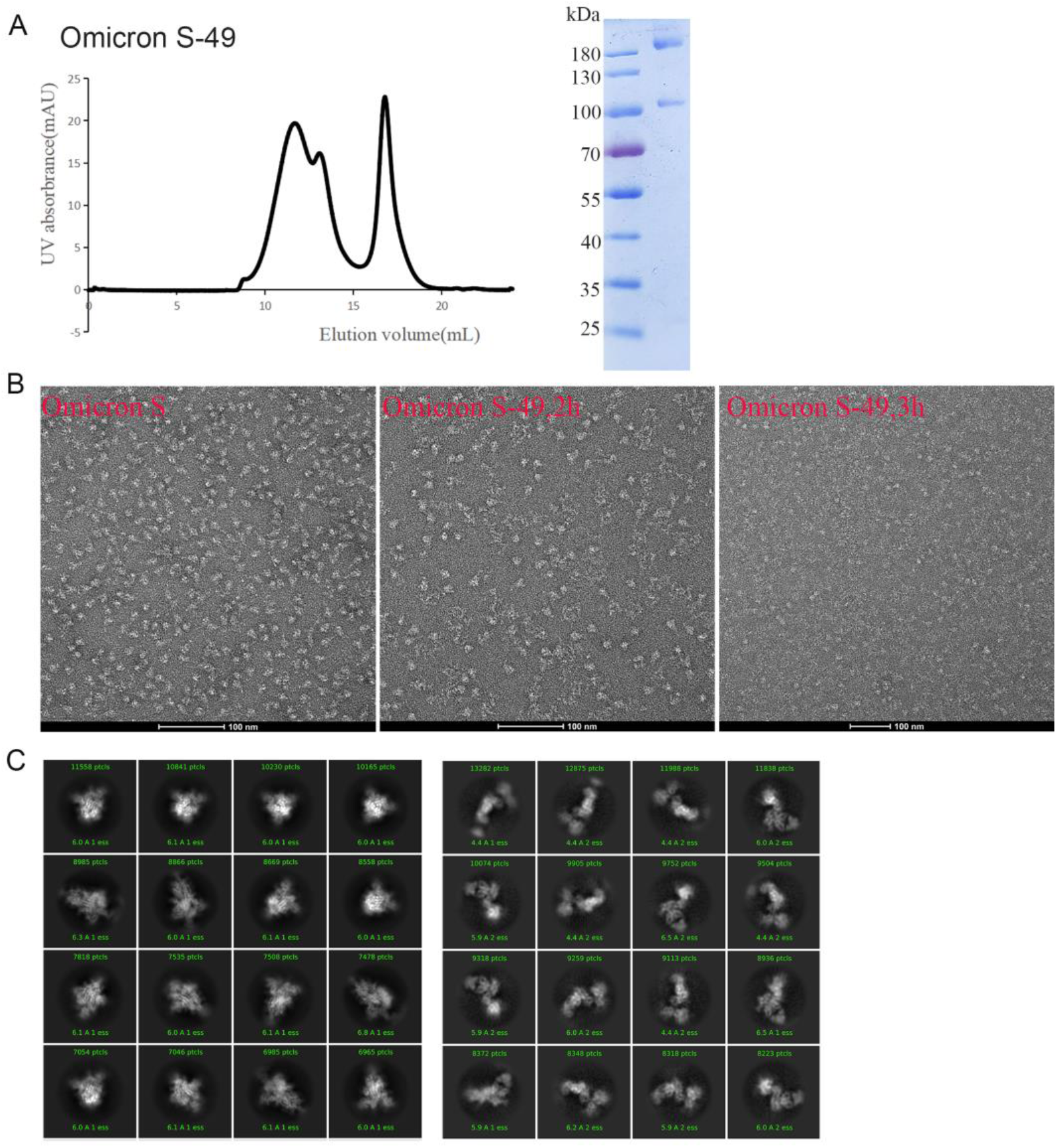
49 binds Omicron S trimer and disassembles it. (A) Purification of 49-Omicron S complex. The gel-filtration curved showed that 49 and S protein can form complex. (B) Negative stain images of 49-Omicron S complex, showing that incubating with 49 for 2h leads to Omicron S trimer disassembling. (C) The 2D classification result of cryo-EM data collected using 1.5h incubation complex, the spike trimer (left) and the spike protomer with 49 (right).

**FIG S3.**
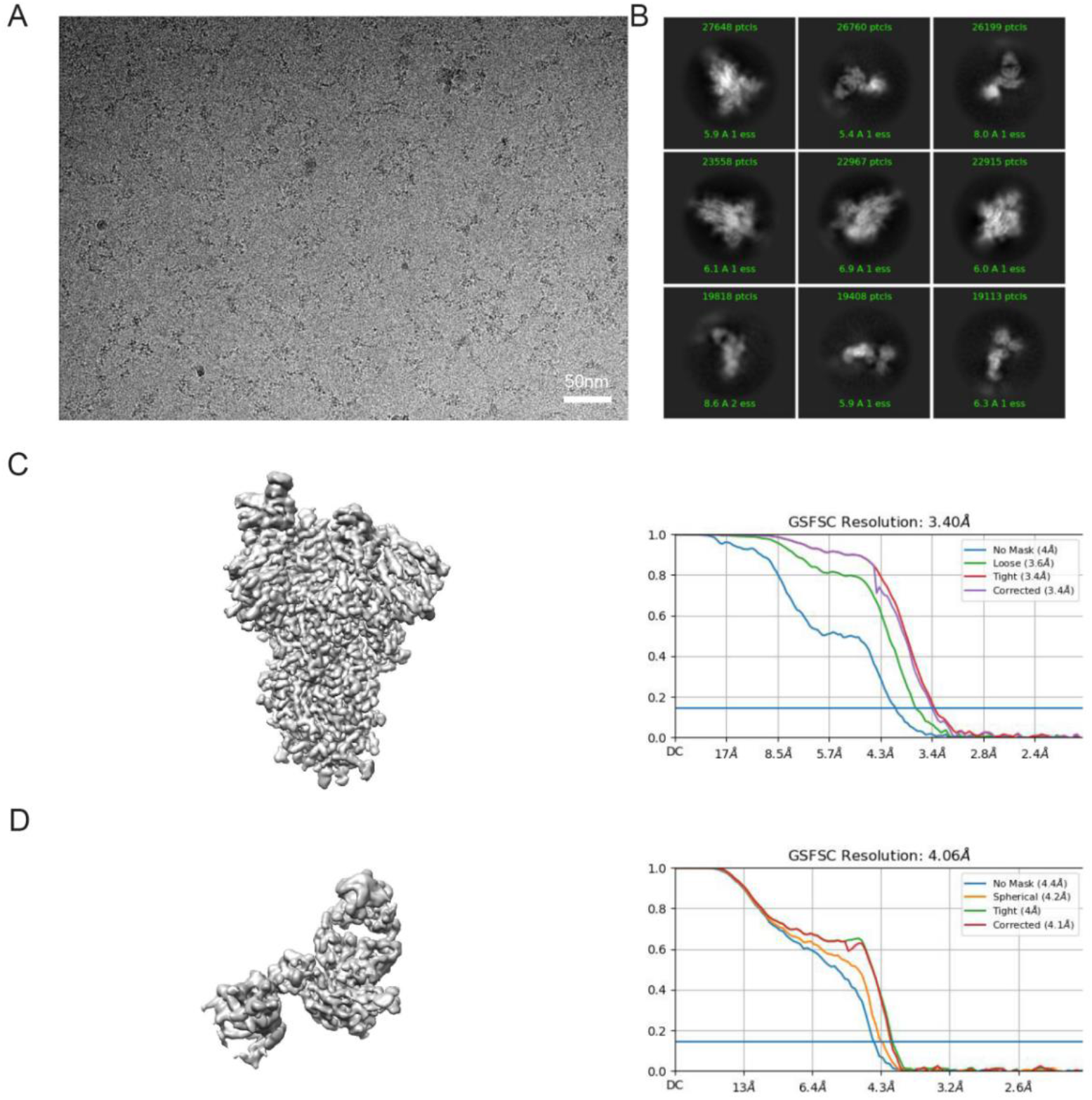
Cryo-EM data collection and processing of SARS-CoV-2 Omicron protomer in complex with 49. (A) Representative electron micrograph of 49 bound SARS-CoV-2 S. (B) 2D classification results of 49 bound SARS-CoV-2 Omicron Spike protein. (C) The reconstruction map of SARS-CoV-2 Omicron Spike trimer and its gold-standard FSC curve from cryoSPARC. (D) The reconstruction map of 49 bound SARS-CoV-2 Omicron Spike protomer and its gold-standard FSC curve from cryoSPARC.

**FIG S4.**
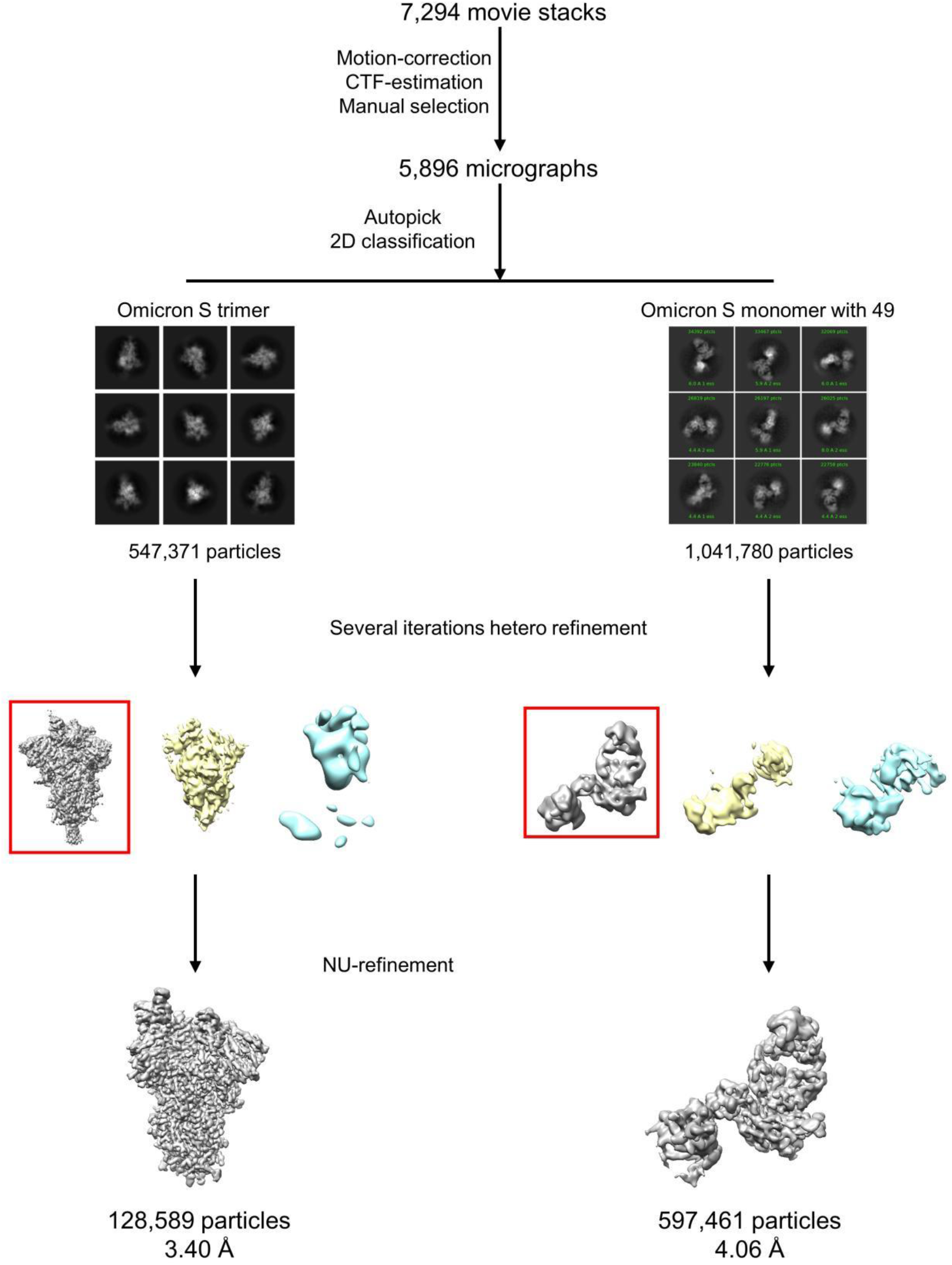
Data processing flowchart of 49 bound SARS-CoV-2 Omicron S protein.

**FIG S5.**
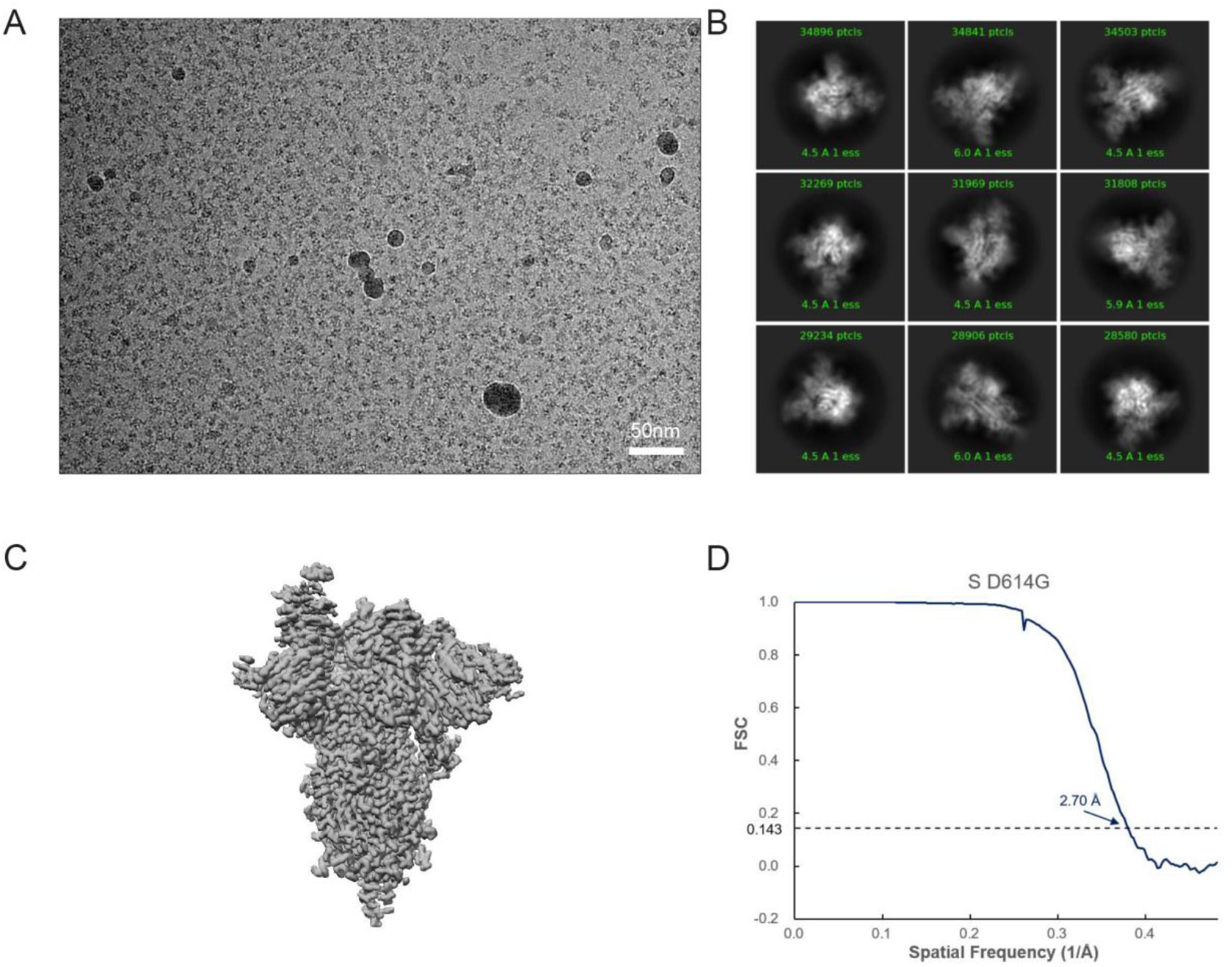
Cryo-EM data collection and processing of S D614G. (A) Representative electron micrograph of SARS-CoV-2 S. (B) 2D classification results of SARS-CoV-2 S. (C) The reconstruction map of SARS-CoV-2 S. (D) Gold-standard Fourier shell correlation curves for each structure. The 0.143 cut-off is indicated by a horizontal dashed line.

**FIG S6.**
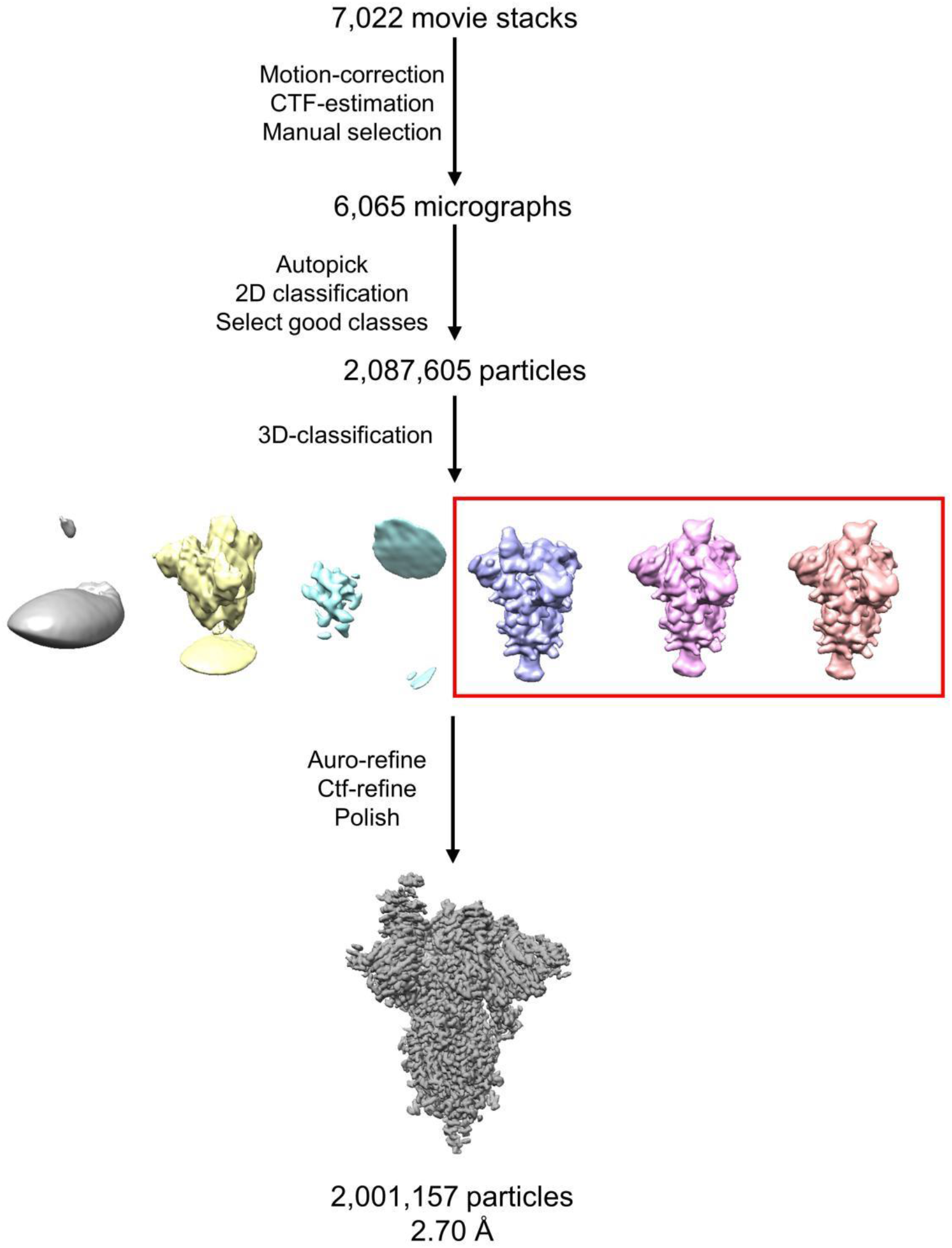
Data processing flowchart of SARS-CoV-2 apo S trimer.

**FIG S7.**
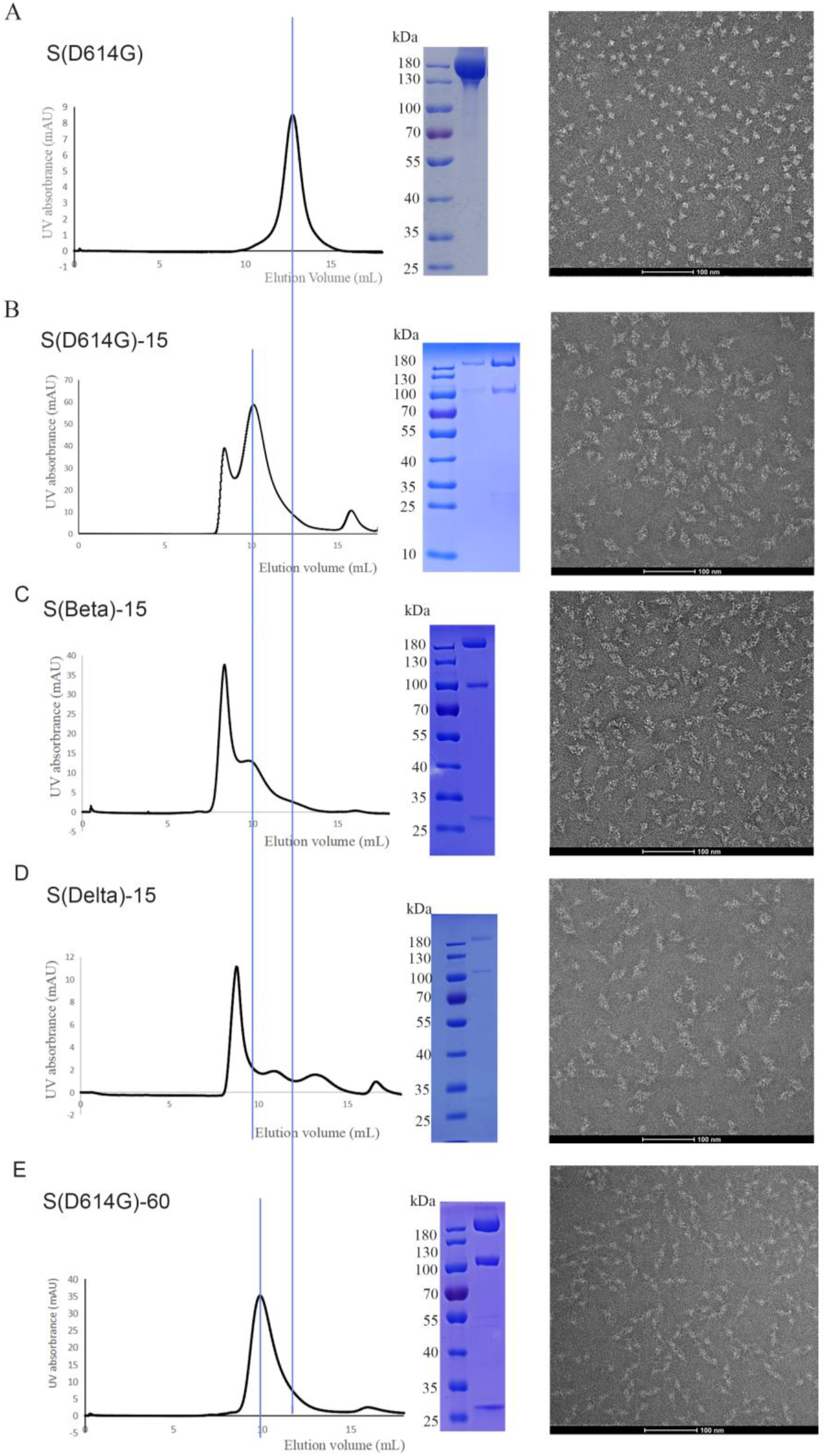
(A-D) Purification and Negative stain images of 15-S (WT, Beta, Delta) complex. The Negative stain image shows that binding of 15 induced Spike trimer forming trimer dimer. (E) Purification and Negative stain images of 60-D614G S complex.

**FIG S8.**
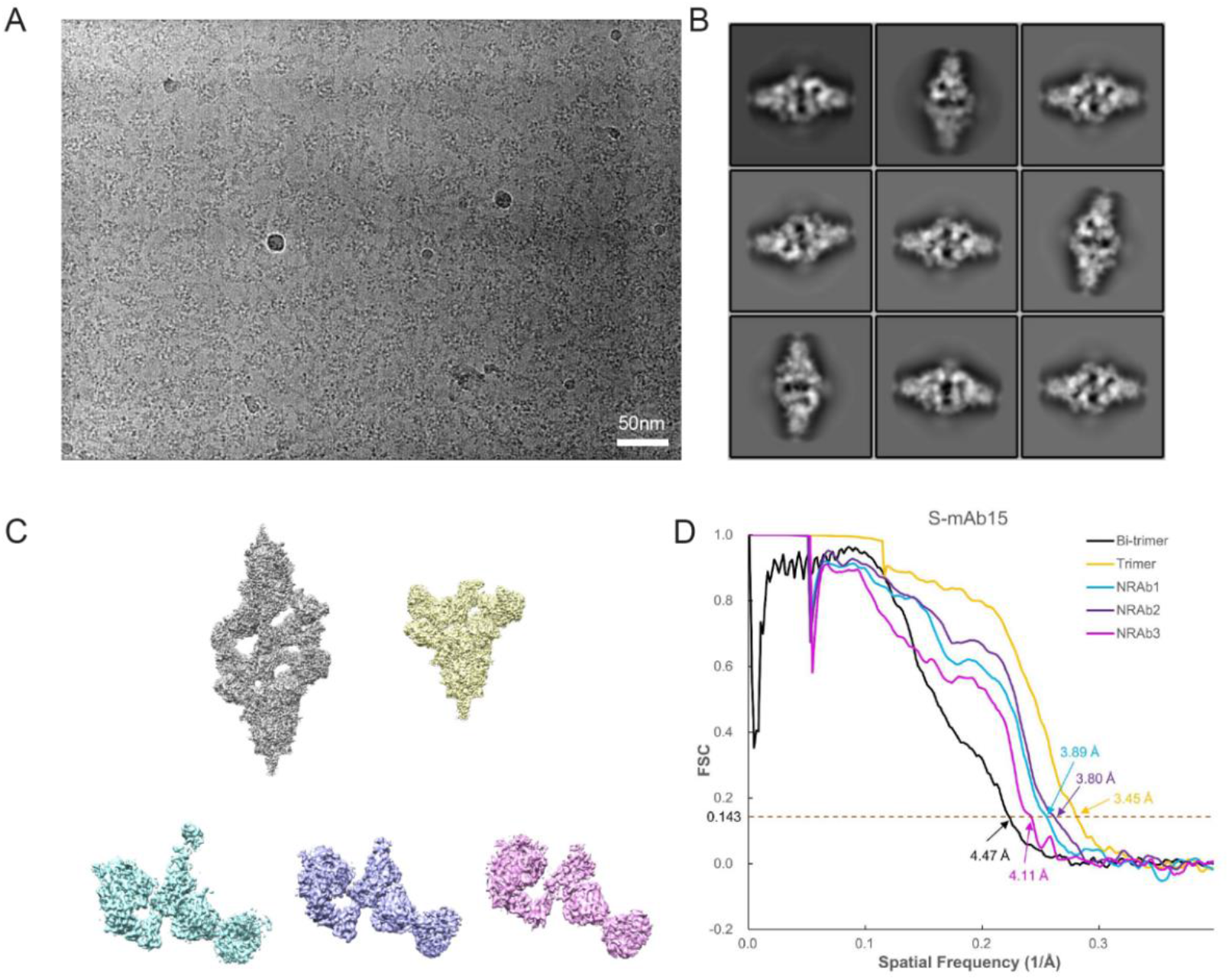
Cryo-EM data collection and processing of 15-S D614G. (A) Representative electron micrograph of 15 bound SARS-CoV-2 S (B) 2D classification results of 15 bound SARS-CoV-2 S. (C) The reconstruction map of trimer dimer, trimer, and three local refinement maps. (D) Gold-standard Fourier shell correlation curves for each structure. The 0.143 cut-off is indicated by a horizontal dashed line.

**FIG S9.**
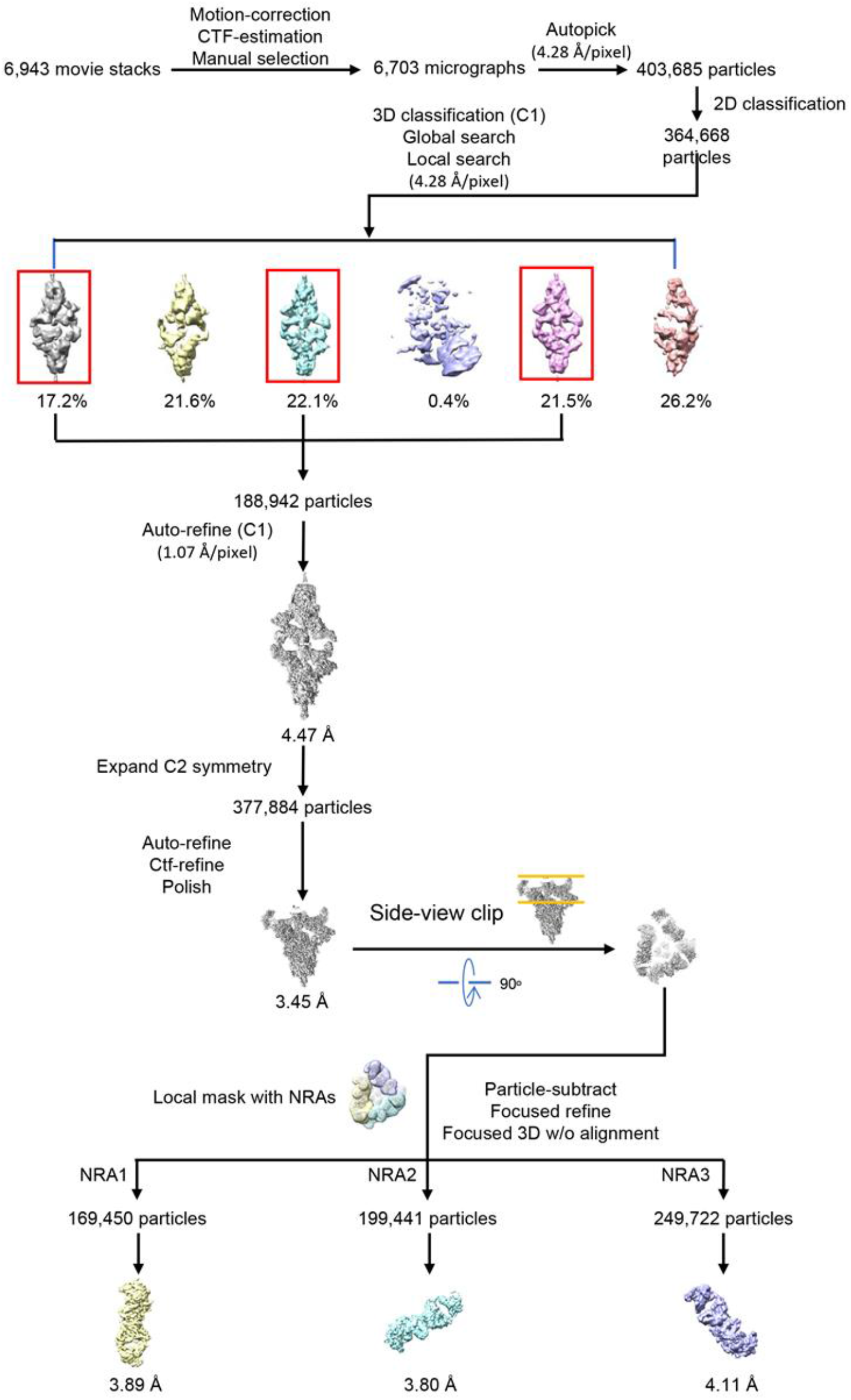
Data processing flowchart of 15 bound SARS-CoV-2 S D614G trimer.

**FIG S10.**
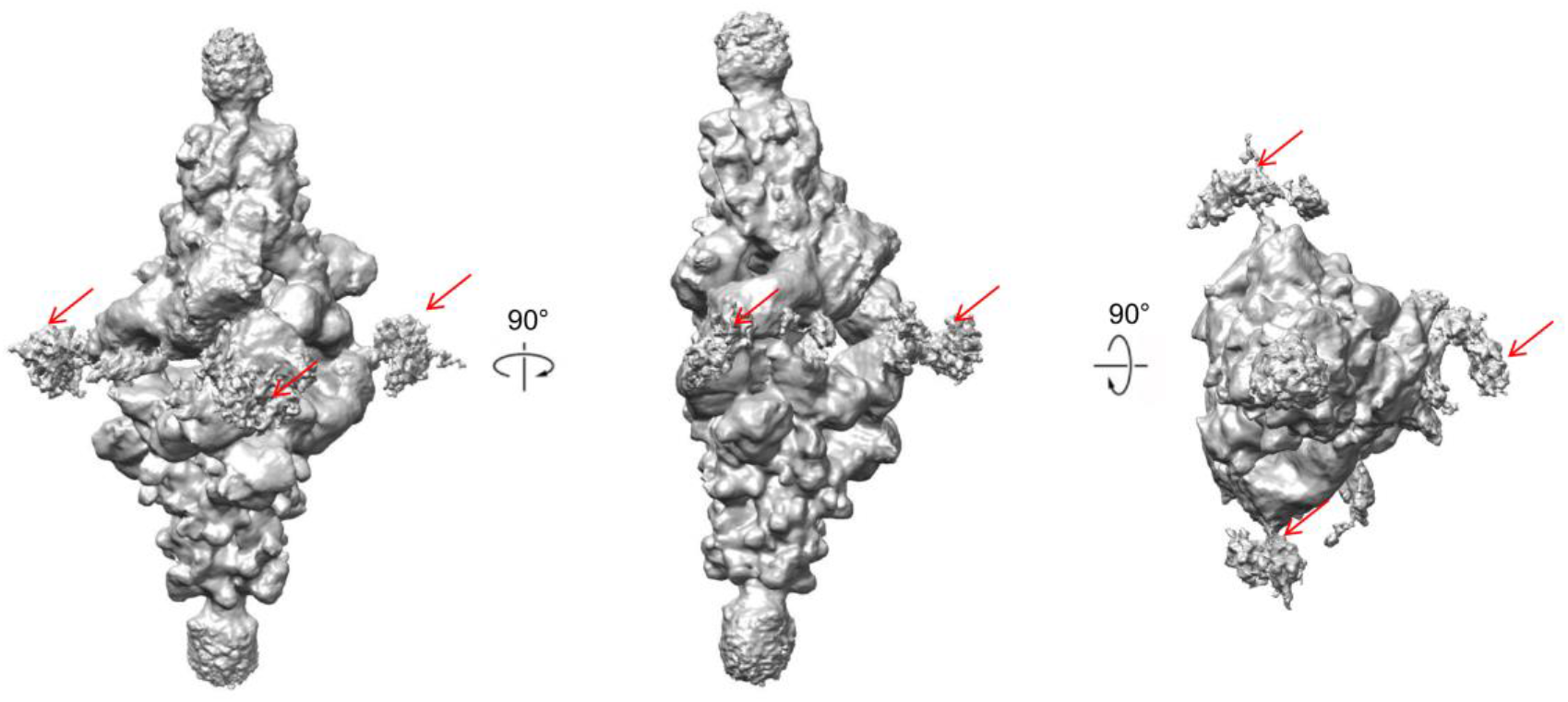
Fc domains of 15-IgGs indicated in cryo-EM data. Fc domain is clear when map is at low range level in Chimera.

**FIG S11.**
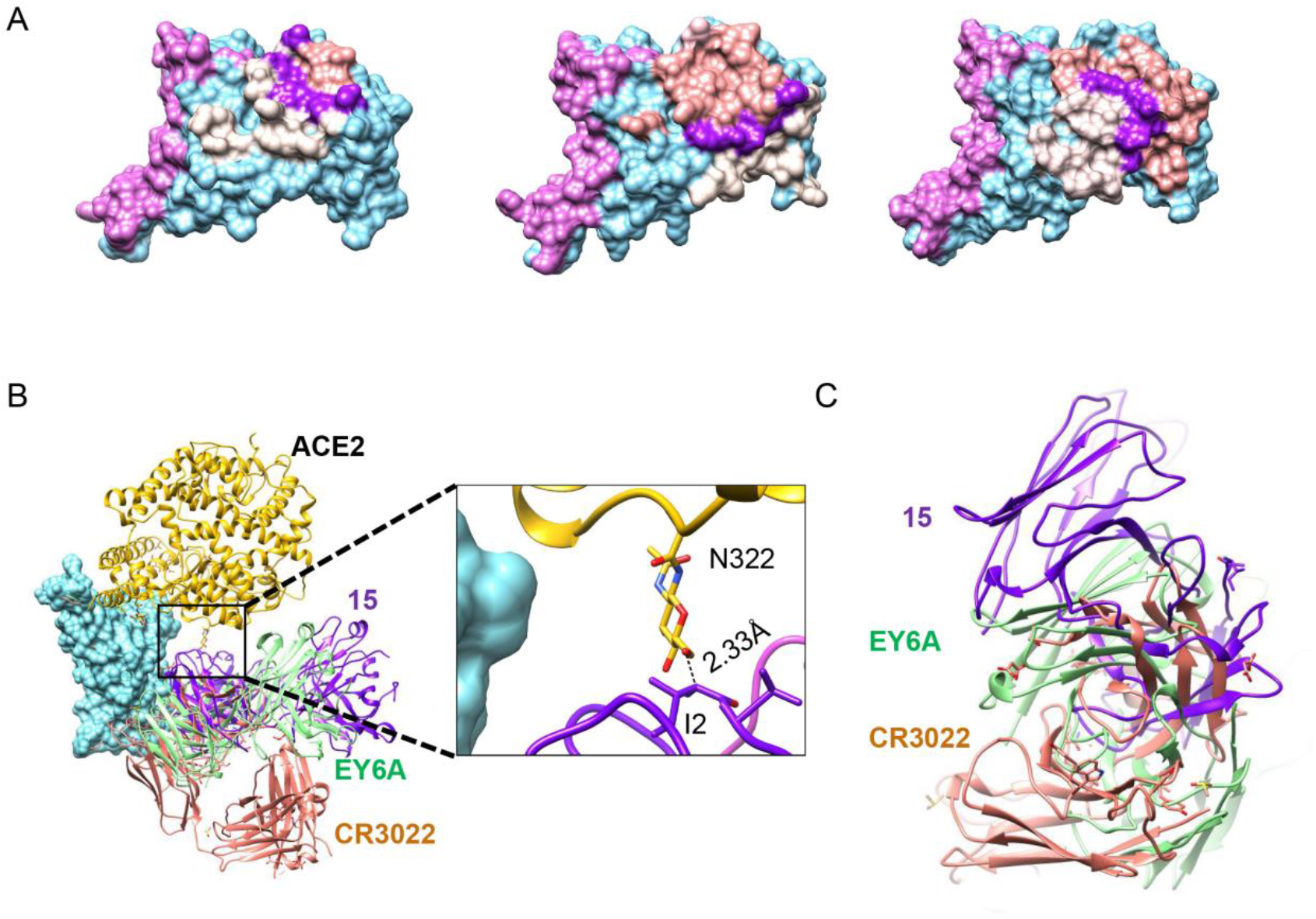
Comparison of 15, ACE2, CR3022, EY6A and their epitopes on the RBDs of SARS-CoV-2. (A) Surface representation of RBD and buried binding site, including 15 (this study), CR3022 (PDB: 6W41), EY6A (PDB: 6ZDG) and ACE2 (PDB: 7T9L). Orchid indicates ACE2 epitope, salmon VH, light gray VL, purple VH and VL, respectively. (B) Comparison of Fab 15, EY6A, CR3022 and ACE2 upon binding to SARS-CoV-2 RBD and 15 would clash with ACE2. (C) The RBD side view of Fab 15, EY6A, CR3022 upon binding to SARS-CoV-2 RBD.

**FIG S12.**
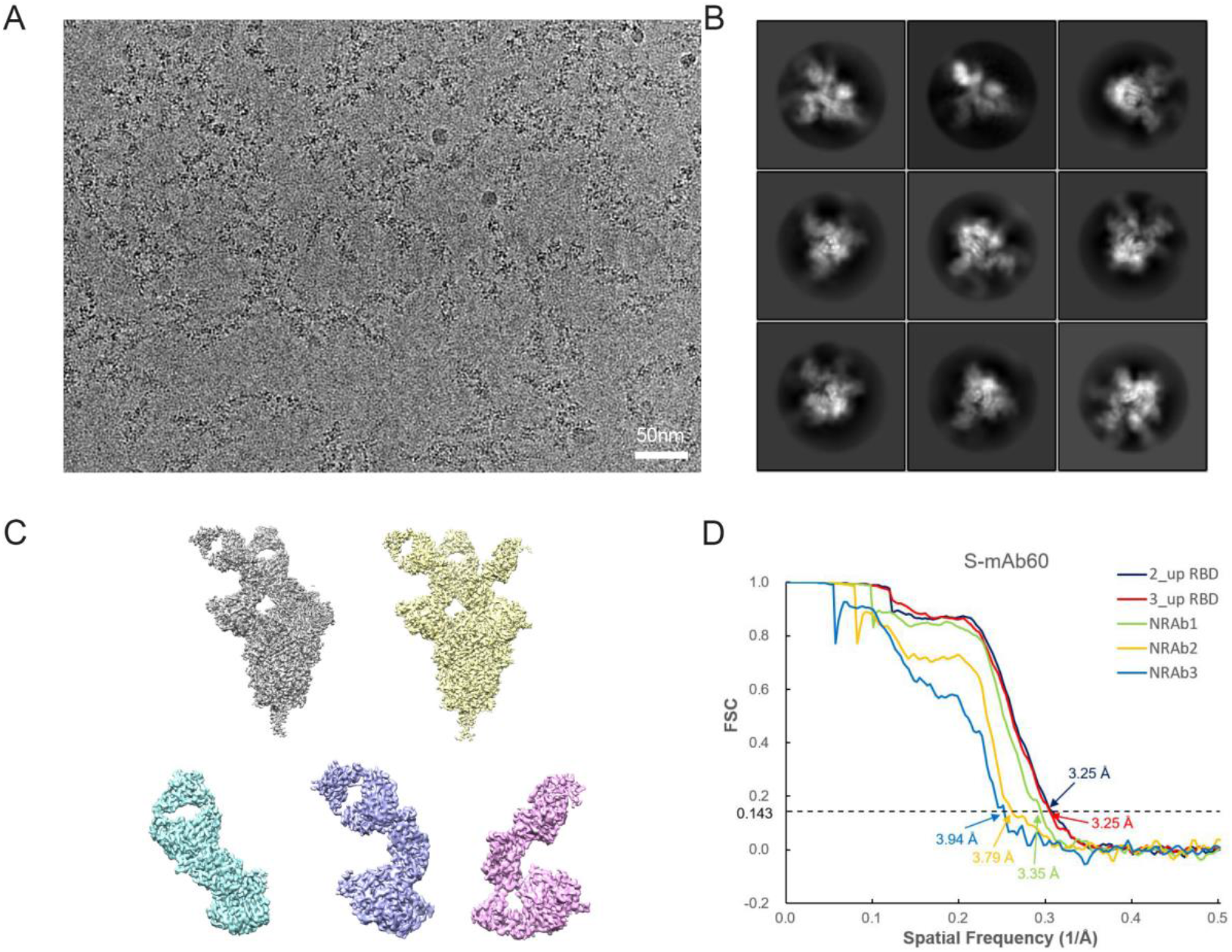
Cryo-EM data collection and processing of 60-S D614G complex. (A) Representative electron micrograph of 60 bound SARS-CoV-2 S. (B) 2D classification results of 60 bound SARS-CoV-2 S. (C) The reconstruction map of 60 bound SARS-CoV-2 S at two states and three locally refined maps. (D) Gold-standard Fourier shell correlation curves for each structure. The 0.143 cut-off is indicated by a horizontal dashed line.

**FIG S13.**
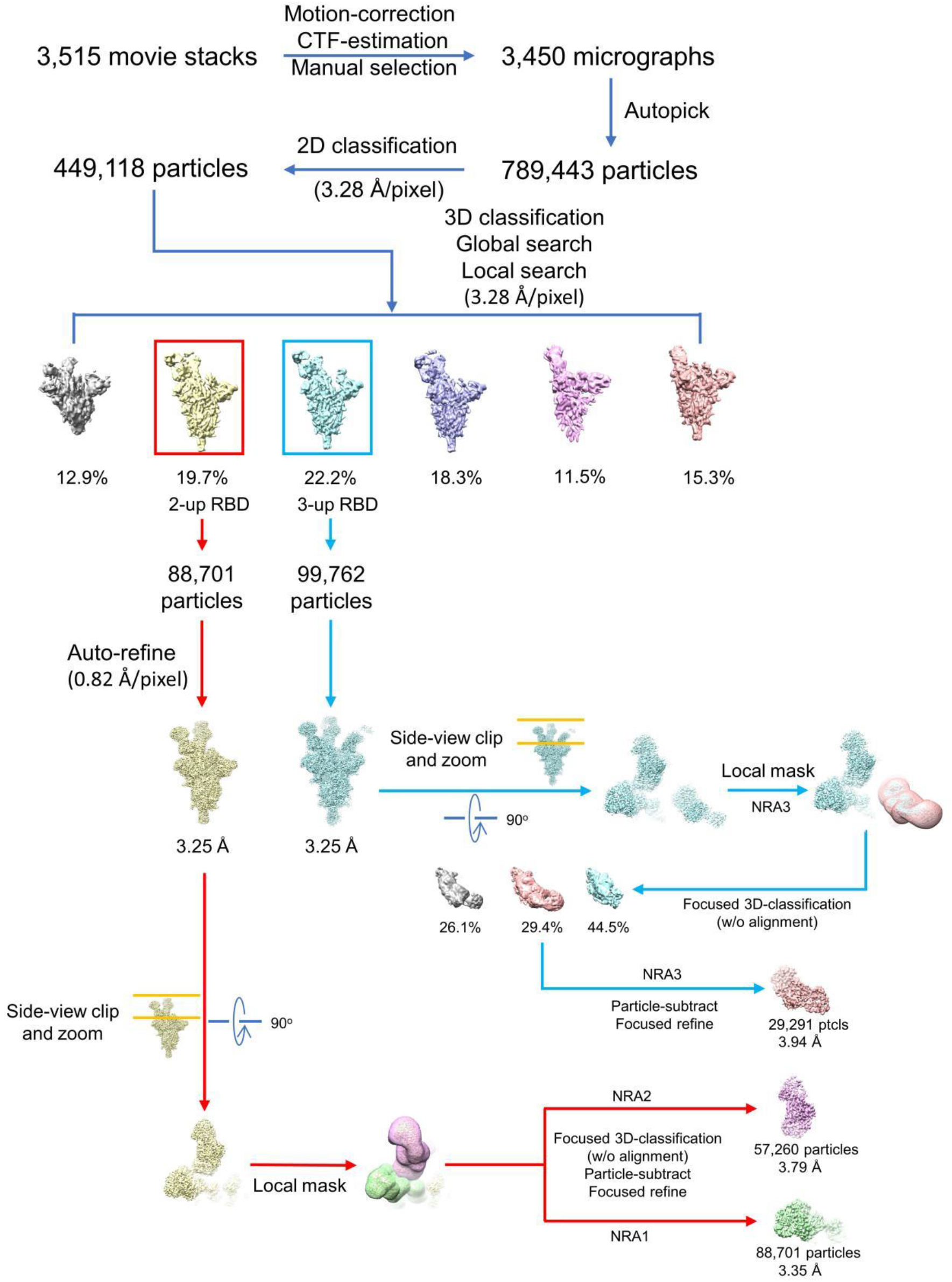
Data processing flowchart of 60 bound SARS-CoV-2 S D614G trimer.

